# Deep learning-guided design of hydrolases for crystalline PET depolymerization

**DOI:** 10.64898/2026.06.04.730138

**Authors:** Banghao Wu, Mingchen Li, Jiaxing Zhang, Jingchuan Li, Xuling Wang, Bozitao Zhong, Jiawei Liu, Yunjie Liu, Bowen Wang, Yang Tan, Wei Qi, Pan Tan, Weishu Zhao, Lirong Zheng, Liang Hong

**Author notes:** These authors contributed equally to this work (Banghao Wu, Mingchen Li, Jiaxing Zhang, Jingchuan Li). These authors jointly supervised this work (Lirong Zheng, Liang Hong).

## Abstract

Poly(ethylene terephthalate) (PET), a ubiquitous polyester used in packaging and textiles, persists in the environment due to its high crystallinity and stability, contributing substantially to global plastic pollution. Enzymatic depolymerization by PET hydrolase (PETase) offers a chemically precise and environmentally sustainable route to convert PET into its monomeric building blocks, enabling recycling. However, practical implementation remains hindered by the rigid, crystalline architecture of PET, which restricts enzyme access and necessitates energy-intensive pretreatment to enable efficient depolymerization. In addition, most PETases achieve only partial depolymerization with low terephthalic acid (TPA) yields and accumulate inhibitory intermediates, while limited thermostability and slow surface kinetics further restrict efficiency. To overcome these barriers, we report VenusPETase, an engineered variant of KbPETase designed using PET-Flow, a state-of-the-art computational framework for PETase engineering. Compared with KbPETase, VenusPETase exhibits a 3.4-fold increase in hydrolytic activity, up to a 27-fold improvement after heat treatment of protein, and a 12 °C enhancement in thermostability. VenusPETase exhibits rapid degradation across a wide crystallinity range (8%–50%) at 50 °C, effectively spanning the entire spectrum of commercial PET products. Moreover, VenusPETase outperformed nine high-performance PETases under their respective optimal conditions and degraded untreated PET substrates across 8%–50% crystallinity, producing TPA as over 95% of the released products with minimal intermediate accumulation. X-ray crystallography and molecular dynamics simulations suggest that dynamic modulation, elevated surface electrostatic potential enhance the interaction of VenusPETase with crystalline PET, thereby lowering the hydrolytic energy barrier and improving catalytic performance. We also demonstrate that untreated, postconsumer-PET from nine different products can all be degraded by VenusPETase. The recovered monomers can be directly repolymerized into virgin-quality PET, demonstrating a closed-loop enzymatic recycling process. In a 100 L bioreactor, VenusPETase completely depolymerizes post-consumer crystalline PET (28% crystallinity) within 24 h under 50 °C. These results establish VenusPETase as a robust biocatalyst that enables efficient, closed-loop recycling of crystalline PET under mild conditions.

## Introduction

Plastics have become indispensable to modern society, supporting global industries ranging from packaging and textiles to electronics and healthcare(*1, 2*). Among them, poly(ethylene terephthalate) (PET) is a petroleum-derived polyester produced at multimillion-tonne scales each year for bottles, fibers, and films, owing to its mechanical strength, chemical resistance, and low manufacturing cost(*3-5*). However, the exceptional stability of its aromatic ester backbone renders PET highly persistent in the environment, leading to its widespread accumulation across terrestrial and marine ecosystems(*6-13*). Growing awareness of this environmental burden has intensified efforts to develop sustainable recycling technologies aligned with circular-economy principles(*14-19*). Thus, enzymatic depolymerization by PET hydrolases (PETases) has emerged as a promising biocatalytic strategy for converting PET into its monomers terephthalic acid (TPA) and ethylene glycol (EG), under mild conditions compared with mechanical and chemical recycling methods, offering a molecularly precise and environmentally benign alternative to conventional thermal or chemical processes(*20-24*).

Enzymatic PET depolymerization has advanced rapidly since the initial discovery of PETase from *Ideonella sakaiensis(25)*. Extensive protein engineering efforts have targeted improvements in catalytic activity and thermostability(*2, 26-29*). Approaches ranging from structure-guided design to directed evolution and machine learning have yielded a growing family of high-performance PETases(*30*). Despite these progresses, enzymatic PET recycling remains constrained by both material and catalytic limitations. The densely packed, crystalline structure of PET restricts enzymatic access to internal ester bonds, necessitating energy-intensive pretreatment to reduce crystallinity(*31, 32*). Even with optimized enzymes, depolymerization is frequently incomplete, yielding limited TPA while accumulating inhibitory intermediates such as MHET(*30, 32, 33*). In addition, limited enzyme robustness and slow surface-erosion kinetics under prolonged or large-scale operation restrict process efficiency.

Recent developments in artificial intelligence (AI)–assisted enzyme engineering provide new opportunities to overcome these bottlenecks by enabling predictive and iterative optimization of protein function(*2, 26*). Current computational approaches for PETase design face the challenge of navigating vast combinatorial sequence space while jointly optimizing catalytic activity and stability, which leads to low design efficiency, limited predictive accuracy and frequent trade-offs between function and robustness(*26, 34, 35*). To address this, we developed PET-Flow, a deep learning-guided framework for PETase engineering that integrates two complementary components: a protein language model (PLM)-based strategy for zero-shot single-site mutant design, and a transfer learning-based optimization model that searches the combinatorial mutation space to maximize activity under a stability constraint. PET-Flow enables data-efficient PETase variant design while substantially reducing the experimental search space.

Using PET-Flow, we engineered KbPETase(*36*) into VenusPETase. Relative to KbPETase, VenusPETase shows a 3.4-fold increase in hydrolytic activity, up to 27-fold higher activity after thermal treatment and a 12 °C gain in thermostability. At 50 °C, VenusPETase depolymerizes PET across a broad crystallinity range (8–50%), including highly crystalline substrates that are typically resistant to enzymatic hydrolysis, thereby encompassing the crystallinity spectrum of major commercial PET products. This broad substrate obviates the need for crystallinity-based waste sorting and specialized pretreatment. X-ray crystallography and molecular dynamics (MD) simulations suggest that elevated surface electrostatic potential enhances the interaction of VenusPETase with PET substrates, particularly highly crystalline PET, thereby lowering the hydrolytic energy barrier and improving catalytic performance. VenusPETase also outperforms nine benchmark PETases(*2, 26-28, 35, 37-40*) under their respective optimal conditions, generating final product mixtures containing more than 95% TPA with minimal intermediate accumulation.The recovered monomers were directly repolymerized into virgin-quality PET, and VenusPETase showed the highest activity in a second depolymerization cycle of the regenerated PET, supporting repeated closed-loop enzymatic recycling. In addition, complete depolymerization of post-consumer crystalline PET (28% crystallinity) is achieved within 24 h at 50 °C in a 100 L bioreactor, highlighting the potential industrial scalability of this system. Together, these results establish PET-Flow as an effective AI-guided PETase engineering framework and VenusPETase as a robust biocatalyst for high-efficiency, high-yield depolymerization of crystalline PET.

## Results

### PET-Flow enables the design of VenusPETase

We first compared ten reported wild-type (WT) PETases(*25, 41-46*) and found that KbPETase(*36*) combined favourable catalytic activity and thermostability (**Supplementary Fig. 1**), making it an attractive starting point for engineering because efficient depolymerization of crystalline PET requires both high hydrolytic efficiency and strong thermal robustness(*16, 20*). Moreover, KbPETase exhibits optimal activity at 50 °C, a relatively mild operating temperature that facilitates crystalline PET depolymerization while reducing the energy input required for industrial processing. To improve the hydrolytic activity and thermostability of KbPETase, we developed PET-Flow, a two-round deep learning-guided framework for engineering PETase that integrates zero-shot mutant-effect prediction with supervised combinational optimization (**Fig. 1a**). Given the absence of prior mutational data for KbPETase, the first round applies zero-shot prediction to nominate promising single-site mutations for experimental characterization; the resulting measurements then train the supervised model that drives higher-order mutant design in the second round.

**Fig. 1.**
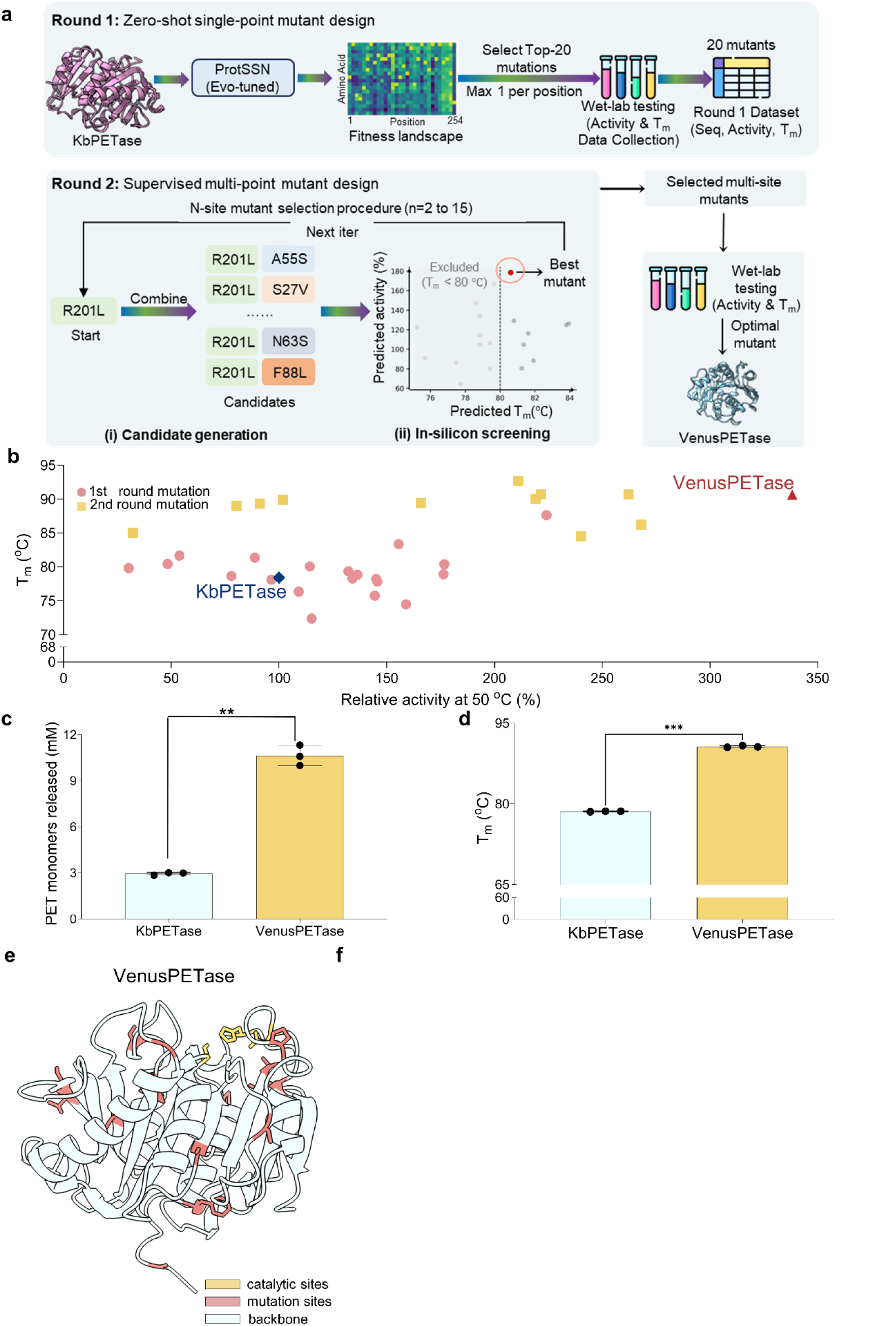
Machine-learning-guided engineering of VenusPETase. **a**, Overview of PET-Flow, an AI-assisted framework for PETase engineering. Because no experimental measurements were initially available for KbPETase mutants, Round 1 began with zero-shot single-site mutant design using ProtSSN-evotuned, which scored all possible single-site mutants and generated a fitness landscape for KbPETase. Then, 20 single-site mutants were selected for experimental measurement of catalytic activity and thermostability (T_m_), thereby establishing the first sequence–activity–stability dataset for KbPETase mutant design. In Round 2, these experimentally determined data were used to train supervised models based on Ankh embeddings and MLP regressors, which were then used to search the combinatorial space of multi-site variants. Candidate higher-order mutants were prioritized by predicted activity under a thermostability constraint (predicted T_m_ ≥ 80 ℃), yielding a focused set of multi-site designs for experimental validation and ultimately the optimized mutant VenusPETase. **b**, Experimentally determined activity–stability landscape of PETase variants generated across three rounds of engineering. Relative activity was measured at 50 °C in 50 mM glycine–NaOH buffer (pH 9.0) using crystalline PET powder (Cry-PET; Goodfellow Cambridge Ltd, Cat. No. ES306000; 39.8% crystallinity) as the substrate, and melting temperature (T_m_) was determined by differential scanning fluorimetry (DSF). Pink circles indicate single-site variants from round 1, whereas yellow squares denote combinatorial variants from round 2. KbPETase and VenusPETase are highlighted in blue and red, respectively. Data are from one independent experiment. **c**, Comparison of the PET-degrading activities of VenusPETase and KbPETase at the optimal reaction temperature of 50 °C. Activity was measured in 50 mM glycine–NaOH buffer (pH 9.0) using Cry-PET as the substrate (mean ± s.d., n = 3, **P < 0.01). T Statistical comparisons were performed using one-sided unpaired t-tests. **d**, Comparison of the thermal stability of VenusPETase and KbPETase. T_m_ was determined by differential scanning fluorimetry (DSF) (mean ± s.d., n = 3, ***P < 0.001). Statistical comparisons were performed using one-sided unpaired t-tests. **e**, Structural distribution of the substitutions incorporated into VenusPETase. The engineered 14 residues (red) and catalytic triad (yellow) are mapped onto the PETase structure to show their spatial distribution across the enzyme scaffold. **f**, Compositional summary of the substitutions incorporated into VenusPETase. The amino-acid replacements retained in the final engineered variant are classified by chemical properties: hydrophobic (gray), polar uncharged (green), special case (pink), and charged (yellow).

In the first round, we used ProtSSN-evotuned, a PETase-specific version of ProtSSN(*47*) fine-tuned on homologous PETase sequences, to predict beneficial single-site substitutions(*48*). This model was selected from PET-Gym, an established benchmark comprising published mutational datasets from multiple PETases for evaluating mutant-effect prediction methods(*26, 35, 38, 49*)(**Supplementary Table 4**), which identified ProtSSN as the best-performing zero-shot predictor and further demonstrated the benefit of homolog-specific fine-tuning (**Extended Data Fig. 1**). To preserve sequence diversity for the downstream optimization, we scored all possible single-site substitutions and selected 20 high-scoring variants spanning distinct residue positions for experimental characterization (**Extended Data Fig. 2a**). Using crystalline PET powder (Cry-PET; Goodfellow Cambridge Ltd, Cat. No. ES306000) with a crystallinity of 39.8% as a model substrate to mimic the crystallinity and post-pretreatment particle size of real post-consumer PET(*31, 33, 35, 50, 51*) and differential scanning fluorimetry (DSF) to assess thermostability(*38, 49*)(**Supplementary Fig. 2**), we found that more than 70% of the tested variants improved catalytic activity or thermostability, and seven variants improved both properties simultaneously(**Fig. 1b**, **Supplementary Table 1**). Given the pervasive trade-off between activity and stability in enzyme engineering(*52-54*), the enrichment of dual-beneficial mutations suggests that PET-Flow effectively identified substitutions capable of partially overcoming this constraint. Among these variants, M3 emerged as the most favorable single-site mutation, increasing PET depolymerization activity by 2.2-fold while raising the melting temperature by 9.2 °C (**Extended Data Fig. 2b**). These results established the first sequence–activity–thermostability landscape for KbPETase and provided the basis for subsequent combinatorial optimization.

In the second round, we trained supervised models using the experimentally measured activities and thermostability of the first-round variants. Guided by PET-Gym benchmarking (**Extended Data Fig. 3**), we selected Ankh(*55*) embeddings coupled with MLP-based regression models to explore the combinatorial sequence space. Candidate variants were evaluated under a thermostability constraint, enabling prioritization of mutants with high predicted activity while excluding unstable designs (**Fig. 1a**). In conventional directed evolution, beneficial mutations often fail to combine productively because of epistatic interactions that diminish activity, stability or both(*56-58*). In contrast, experimental characterization of 12 PET-Flow-designed variants showed that all exhibited improved thermostability, and eight additionally showed enhanced catalytic activity (**Fig. 1b**, **Supplementary Table 2**). Compared with the single-site variants identified in the first round, these higher-order mutants displayed further gains in both properties, indicating that PET-Flow successfully navigated epistatic constraints while requiring only a limited number of experimental measurements(*27, 28, 49*) (**Extended Data Fig. 2c** and **d**). Among these designs, M31 exhibited the best overall performance and was designated VenusPETase (**Extended Data Fig. 2b**).

VenusPETase displayed a melting temperature of 90.7 °C, representing a 12 °C relative to KbPETase (**Fig. 1d**). This substantial thermostability enhancement was achieved without introducing additional disulfide bonds (**Fig. 1e** and **f**), in contrast to many previous PETase-engineering strategies(*28, 35, 37, 38*). VenusPETase also showed substantially improved depolymerization activity on crystalline PET across a broad temperature range, with 3.4-, 14- and 22-fold higher activity than KbPETase at 50, 60 and 65 °C, respectively (**Fig. 1c**, **Extended Data Fig. 4b**). Kinetic analysis(*25, 36, 49, 59*) further demonstrated enhanced catalytic efficiency relative to KbPETase and FastPETase, which also has an optimum temperature of 50 °C (**Extended Data Fig. 4c**, **Extended Data Table. 1**). Because TPA is the preferred product for downstream recycling, product distribution is a critical determinant of process economics(*20, 33, 60, 61*). VenusPETase increased the TPA fraction of the final hydrolysis products to 96.09% (**Extended Data Fig. 4a**), substantially reducing accumulation of intermediate species and potentially lowering purification costs. Enhanced robustness was further demonstrated by heat-inactivation experiments, in which VenusPETase retained substantial activity after incubation at 65 °C for 72 h, whereas KbPETase lost activity completely (**Extended Data Fig. 5**). Moreover, VenusPETase maintained high activity across a broad crystallinity range, from amorphous to highly crystalline materials, and exhibited stronger degradation activity toward crystalline than amorphous PET under the tested conditions. By contrast, the activity of KbPETase decreased with increasing substrate crystallinity (**Extended Data Fig. 4d**), highlighting the suitability of VenusPETase for heterogeneous post-consumer PET. Together, these results demonstrate that PET-Flow enables simultaneous optimization of catalytic activity, thermostability and product selectivity within a single PETase scaffold, yielding VenusPETase as a highly efficient catalyst for crystalline PET depolymerization.

### Structural and dynamic basis for the enhanced catalytic efficiency of VenusPETase on PET

To elucidate the molecular basis for the enhanced catalytic activity of VenusPETase on substrates of varying crystallinity, we performed X-ray crystallography and molecular dynamics simulations. Structural alignment showed that VenusPETase (PDB ID: 9VJ7) adopts a canonical *α/β*-hydrolase fold nearly identical to that of KbPETase (PDB ID: 9IW9) (RMSD = 0.281 Å), indicating that the overall protein architecture was conserved after engineering (**Supplementary Fig. 3**). Based on the crystal structures of KbPETase and VenusPETase, we conducted a two-stage MD study of enzyme–substrate interactions. First, we constructed PET substrate models at high (CPET) and low (APET) crystallinity, respectively (**Supplementary Fig. 4** and **5**), and investigated their interactions with both KbPETase and VenusPETase (**Fig. 2a**). After 50 ns MD simulation, VenusPETase induced stronger disruption of CPET structure than KbPETase, resulting in a concave curvature on CPET surface (**Fig. 2b**). This could be due to the increased electrostatic surface potential of VenusPETase (**Supplementary Fig. 6**), reflected in an enhancement in solvation energy. This suggests that VenusPETase maintains a structural advantage that facilitates increased contacts with CPET through enhanced surface charge, potentially contributing to more efficient CPET degradation. Interaction energy analysis further revealed that VenusPETase exhibits a lower average interaction energy with CPET (−482.524 kJ/mol) compared to KbPETase (−332.183 kJ/mol), reflecting its stronger binding affinity with CPET. Notably, the average interaction energy of VenusPETase and KbPETase with APET were −436.206 and −335.737 kJ/mol, respectively, indicating a general enhancement in VenusPETase’s affinity for PET substrates across different crystallinities, consistent with experimental observations (**Supplementary Fig. 7**, **Extended Data Fig. 4d**). The enhanced interaction of VenusPETase with CPET (more negative interaction energy) correlates directly with the ability to overcome the high kinetic and thermodynamic barriers associated with crystalline surface binding. Thus, the enhanced interaction of VenusPETase was especially consequential for highly crystalline PET, for which efficient surface engagement is typically significant to PET enzymatic depolymerization. Collectively, VenusPETase achieves its advantage on high-crystallinity PET through electrostatic surface optimization that promotes substrate distortion and tighter binding, thereby enhancing crystalline PET degradation, for which efficient surface engagement is typically a key barrier to PET enzymatic depolymerization(*62, 63*).

**Fig. 2.**
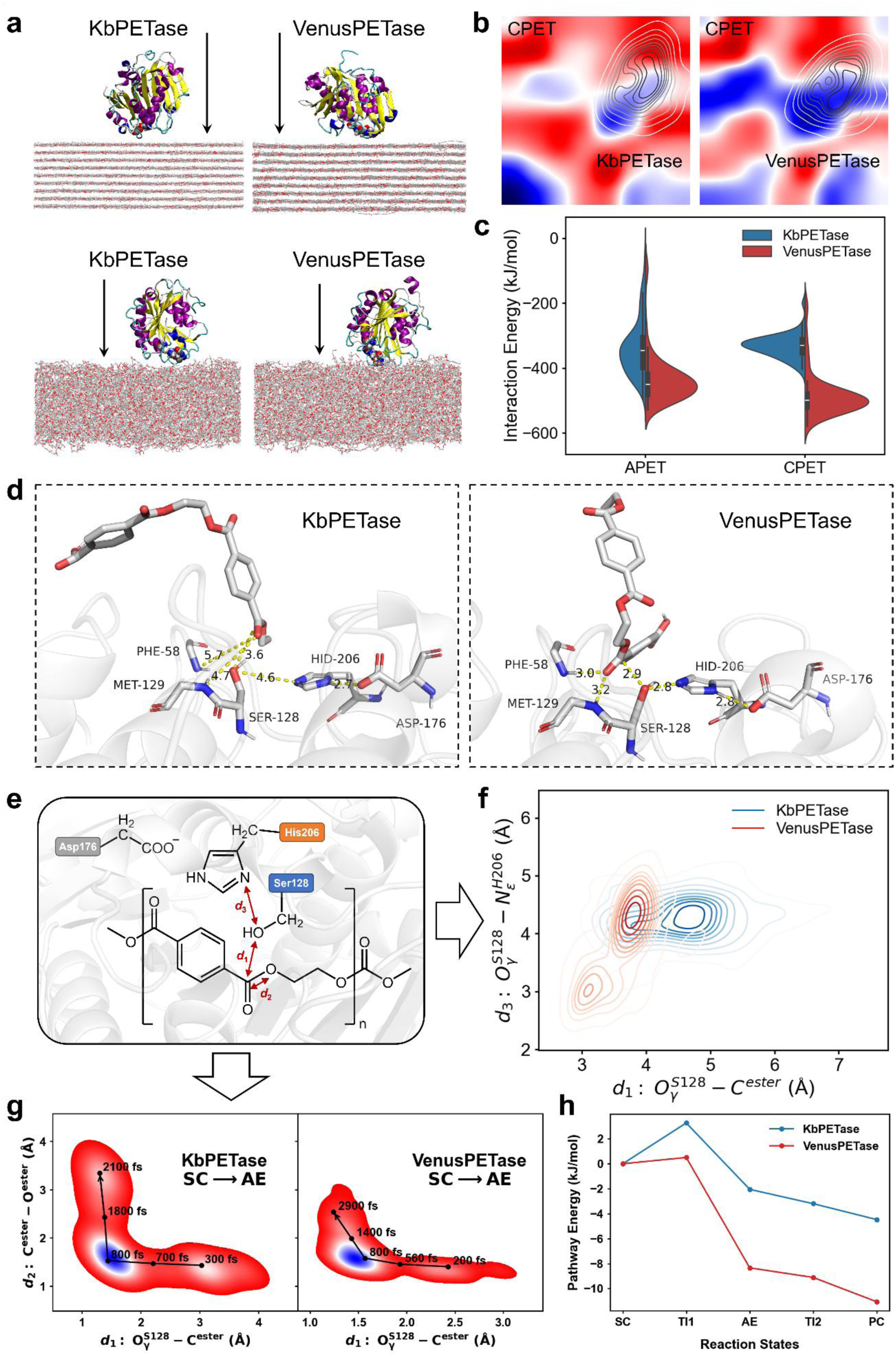
Interaction with amorphous and crystalline PET substrates of VenusPETase and KbPETase. **a**, Molecular dynamics conformations of KbPETase and VenusPETase with high-crystallinity CPET (first row) and low-crystallinity APET (second row) after 100 ns MD simulations. **b**, Curvature analysis of CPET after interaction with VenusPETase. The red areas indicate the convexities of the PET film, and the blue areas correspond to the concavities. The inset contour reveals the position of both enzymes, where VenusPETase results in a larger concave area compared to KbPETase. **c**, Binding energy of both enzymes to the APET and CPET substrates. VenusPETase shows stronger binding energy with both substrates than KbPETase. The binding affinity of VenusPETase with CPET is higher than with APET, which is consistent with the experiments. **d**, Conformations of KbPETase and VenusPETase with 2PET model substrates after 50 ns MD simulations. The KbPETase did not maintain effective catalytic conformations, while VenusPETase shows effective catalytic structure of substrate and catalytic triad, with close key distances (yellow lines). **e**, The key distances related to the PET degradation process (annotated by red arrows): *d*_1_, catalytic distance (O^S128^—C^ester^); *d*_2_, breaking bond length (O^ester^—C^ester^); *d*_3_, electron transfer distance (O^S128^—N^H206^). **f**, The two-dimensional distribution of *d*_1_ and *d*_3_ among metadynamics trajectory. VenusPETase showed a prominent distribution *d*_1_ = 3 Å and *d*_3_ = 3 Å, indicating a more active structural conformation compared to KbPETase. **g**, Estimated free energy landscape of *d*_1_ and *d*_2_ during meta-FPMD simulation. Left panel: SC to TI reaction step of KbPETase. Right panel: SC to TI reaction step of VenusPETase. **h**, Estimated energy of the reaction states. Among meta-FPMD simulations, VenusPETase showed smaller energy barriers of TI state than KbPETase during the reaction process, indicating higher catalytic ability of PET degradation.

To further compare the catalytic mechanisms of the two enzymes, we also performed both force field-based MD and first-principles MD (FPMD) analysis on a 2PET model substrate. Regarding classic MD, throughout 50-ns simulations, the 2PET substrate adopted an unstable conformation in complex with KbPETase, deviating from the catalytically productive pose (**Fig. 2d**). The distances between the ester carbonyl oxygen of 2PET and the oxyanion hole residues F58/M129 were 5.3 Å and 4.7 Å, which is too far to support effective transition-state stabilization. In contrast, VenusPETase maintained a catalytically competent conformation, with key oxyanion hole distances of 3.0 Å and 3.2 Å. Other key catalytic distances, such as O_γ_^S128^ – C^ester^ and O_γ_^S128^ – N_ɛ_^H206^, were consistently below 3 Å, underscoring VenusPETase’s superior capacity to orient the substrate into a reactive configuration. This phenomenon was further corroborated by metadynamics simulations (**Fig. 2e**), which revealed a free energy minimum near 3 Å along two collective variables (CVs) for the VenusPETase/2PET system, consistent with an optimal catalytic distance. However, no such minimum was observed for KbPETase/2PET.

Meanwhile, regarding FPMD simulations, we studied the enzyme-substrate complex structures after MD equilibration (**Fig. 2e** and **f**). First, according to the reaction mechanism related to the catalytic triad (S128, D176, H206) in PET hydrolases (**Extended Data Fig. 6a**), we focused on the five key states of the reaction: substrate-enzyme complex (SC), first tetrahedral intermediate (TI1), acyl-enzyme (AE) intermediate, second tetrahedral intermediate (TI2), and product complex (PC). To monitor ester bond cleavage and new bond formation, we defined two collective variables as the distance of O_γ_^Ser128^ – C^ester^ (*d*_1_) and C^ester^ – O^ester/water^ (*d*_2_) (**Fig. 2e**). The free energy surface of (*d*_1_, *d*_2_) revealed differences between KbPETase and VenusPETase during two key steps: acylation (SC→TI→AE) and deacylation (AE→TI→PC) (**Fig. 2e**, **Extended Data Fig. 6b**). In both systems, the 2PET substrate underwent the full reaction pathway, and finally converted to products. Meanwhile, the energy obtained by kernel density estimation (**Fig. 2f**) among the pathways showed that VenusPETase exhibited a smaller energy barrier than KbPETase, which speeds up the rate-limitation SC→TI→AE step, and finally enhanced the thorough catalytic performance of VenusPETase (The detailed reaction snapshots and processes were illustrated in **Extended Data Fig. 6c** and **Video S1-4** in **Supplementary file 2**).

### VenusPETase outperforms state-of-the-art PET hydrolases

Having established the structural and dynamic basis for the enhanced catalytic performance of VenusPETase, we next asked whether these molecular advantages translate into functional superiority under practical and industrially relevant conditions. Therefore, we benchmarked VenusPETase against a panel of leading engineered PETases, including TurboPETase(*49*), FastPETase(*2*), LCC^ICCG^(*37*), DepoPETase(*27*), HotPETase(*28*), PES-H1^L92F/Q94^(*39*), ThermoPETase(*40*), CaPETase^M9^(*35*) and Kubu-P^M12^(*38*) (**Fig. 3**). Across all enzymes and reaction temperatures tested, VenusPETase showed the highest PET depolymerization activity (**Fig. 3a**). At 50 °C, VenusPETase displayed 1.9-fold higher activity than LCC^ICCG^ and TurboPETase at 65 °C, and 2–4-fold higher activity than PETases optimized for 50 °C, including Kubu-P^M12^ and FastPETase. Interestingly, these gains were achieved despite operating below the glass transition temperature of PET(*64, 65*), a regime in which efficient enzymatic depolymerization remains challenging. VenusPETase also showed a broad catalytic temperature profile. Its activity at 60 °C exceeded that of all comparator enzymes across the tested temperature range and remained 1.8–4-fold higher than those of thermostable PET hydrolases optimized for 60 °C, including CaPETase^M9^ and HotPETase. Furthermore, whereas most PETases optimized for 50 °C exhibited little or no detectable activity at 65 °C, VenusPETase retained substantial catalytic activity under these conditions (**Fig. 3a**).

**Fig. 3.**
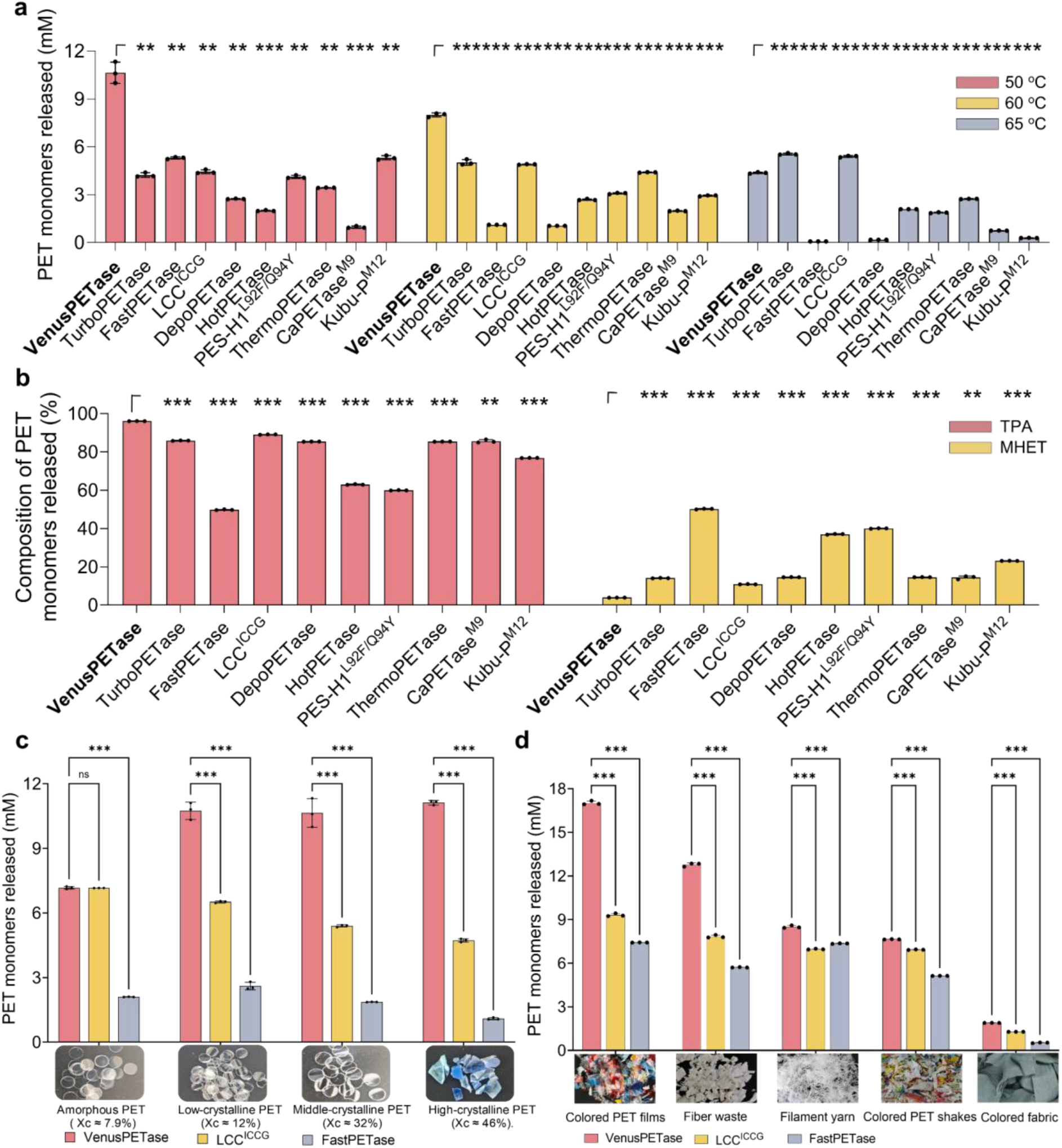
VenusPETase exhibits superior catalytic activity and TPA yield across PET substrates with varying crystallinity levels and across post-consumer PET. **a**, Comparison of PET-hydrolytic activity of VenusPETase and nine benchmark PET hydrolases towards crystalline PET powder at three temperatures: 50 °C (red), 60 °C (yellow), and 65 °C (blue). Reactions were performed at an enzyme loading of 3 mg_enzyme_ g_PET_^−1^ in respective optimal buffer for each enzyme.All data represent mean ± s.d. from three independent biological replicates (n = 3). Statistical analysis was performed using two-way ANOVA followed by Dunnett’s multiple comparisons test, comparing the other two enzymes with my enzyme within each substrate **P < 0.01, ***P < 0.001; n.s., not significant. **b**, Comparison of the product distribution generated by VenusPETase and benchmark PET hydrolases during PET depolymerization. Reaction products generated under the respective optimal temperature and buffer conditions for each enzyme were analysed by UPLC. Red bars denote the proportion of TPA, and yellow bars denote the proportion of MHET. Statistical comparisons were performed using one-sided unpaired t-tests. **P < 0.01; ***P < 0.001; n.s., not significant. **c**, Depolymerization of PET substrates with different crystallinities by VenusPETase and FastPETase at 50 °C, and by LCC^ICCG^ at 65 °C. Representative photographs of PET substrates used in crystallinity-dependent assays: amorphous PET (AF-PET, Goodfellow Cambridge Ltd, Cat. No. ES301445; X_c_ ≈ 7.9%), low-crystalline PET (disposable fruit containers; X_c_ ≈ 12%), middle-crystalline PET (mineral water bottle walls; X_c_ ≈ 32%), and high-crystalline PET (commercially recycled colored PET flakes; X_c_ ≈ 46%). All substrates were pretreated with liquid nitrogen and cryoground into 150 μm powders for degradation assays. **d**, Bar plots compare the degradation efficiencies of VenusPETase (red), LCC ICCG (yellow), and FastPETase (gray) across five distinct PET materials, including colored waste PET films, mixed-color PET flakes, white polyester fiber production offcuts, polyester filaments, and colored waste polyester fabric scraps. Reactions were conducted under the respective optimal catalytic temperature and buffer conditions of each enzyme. Representative images of the tested PET materials are shown below. All data represent mean ± s.d. from three independent biological replicates (n = 3). Statistical analysis was performed using two-way ANOVA followed by Dunnett’s multiple comparisons test, comparing the other two enzymes with my enzyme within each substrate **P < 0.01, ***P < 0.001 (**d,e**).

This broad temperature adaptability is consistent with its enhanced thermostability, as DSF revealed a melting temperature of 90.7 °C for VenusPETase (**Extended Data Fig. 7**, **Supplementary Fig. 8**). To further assess operational robustness, we compared VenusPETase with FastPETase, which shares the same optimal reaction temperature, and LCC^ICCG^, a leading thermostable benchmark enzyme. To assess long-term thermal robustness(*27, 35*), VenusPETase, FastPETase, and LCC^ICCG^ were pre-incubated for 72 h at both 50 °C and 65 °C, representing the optimal catalytic temperatures of FastPETase/VenusPETase and LCC^ICCG^, respectively. After heat treatment, VenusPETase retained 63% and 51% of its initial activity after pre-incubation at 50 °C and 65 °C. In both residual activity and absolute PET degradation activity after heat treatment, VenusPETase outperformed FastPETase and LCC^ICCG^, with residual activities 2.3–25 fold higher than those of the comparator enzymes (**Extended Data Fig. 5**). These results demonstrate not only enhanced thermostability but also prolonged operational durability(*21, 50, 66*), a key determinant of industrial process economics.

Because complete conversion of PET products into TPA is preferred for downstream chemical recycling and repolymerization(*20, 33, 50*), we next compared product distributions among the tested PETase (**Fig. 3b**). Under their respective optimal reaction conditions, VenusPETase converted approximately 95% of released PET-derived monomers into TPA, whereas all other PETases accumulated more than 10% mono(2-hydroxyethyl) terephthalate (MHET) (**Fig. 3b**). This high TPA selectivity minimizes intermediate accumulation and simplifies downstream monomer recovery, thereby improving the suitability of VenusPETase for closed-loop recycling applications. In addition, these results also demonstrate VenusPETase as a leading PETase across multiple performance metrics (**Extended Data Fig. 8**).

We next evaluated whether the superior catalytic performance of VenusPETase extends to PET substrates with different crystallinities. VenusPETase was benchmarked against LCC^ICCG^ and FastPETase using representative PET materials spanning the crystallinity range commonly encountered in commercial and post-consumer products, including commercial amorphous PET (AF-PET, Goodfellow Cambridge Ltd, Cat. No. ES301445; X_c_ ≈ 7.9%), disposable fruit containers (low-crystalline PET; X_c_ ≈ 12%), water bottles (middle-crystalline PET; X_c_ ≈ 32%), and colored PET pellets (high-crystalline PET; X_c_ ≈ 46%) (**Supplementary Fig. 9**, **Extended Data Table 2**). VenusPETase maintained high activity across all substrates and was the only enzyme that exhibited a preference for crystalline PET over amorphous PET. Across crystalline PET substrates, VenusPETase showed 1.6–10-fold higher depolymerization activity on crystalline PET substrates than the comparator enzymes (**Fig. 3c**). VenusPETase also exhibited the highest activity toward colored PET, a challenging substrate class in which pigments and additives often impede enzymatic hydrolysis(*35, 62*).

To assess performance on realistic waste streams, we further evaluated VenusePETase using diverse post-consumer PET products(*1*). Five representative PET waste materials were treated with VenusPETase at 50 °C, resulting in substantially higher monomer release than that achieved by LCC^ICCG^ or FastPETase across all products tested (**Fig. 3d**). A multidimensional comparison incorporating catalytic activity, thermostability and product selectivity further identified VenusPETase as the top-performing enzyme across all evaluated metrics (**Extended Data Fig. 9**). These results establish VenusPETase as a leading PET hydrolase that combines exceptional catalytic efficiency, broad crystallinity tolerance, high TPA selectivity and operational robustness, enabling efficient depolymerization of diverse post-consumer PET feedstocks.

### Complete depolymerization of post-consumer PET waste by VenusPETase

To evaluate the applicability of VenusPETase for closed-loop PET recycling, we first tested its ability to depolymerize intact post-consumer PET products at 50 °C (**Fig. 4a**). An untreated disposable fruit container as the substrate (12 cm × 16 cm × 0.5 cm, crystallinity gradient 4.8–12.7%) (**Supplementary Fig. 10**, **Extended Data Table. 2**). At 50 °C, VenusPETase depolymerized the entire container within 30 hours. Degradation was first observed in more crystalline sidewall regions and subsequently progressed to the less crystalline bottom region (**Fig. 4a**, left panel; **Video S5**), consistent with the preferential activity of VenusPETase toward crystallinity PET substrates (**Fig. 3c**). To demonstrate close-loop recycling, the released monomers, TPA and EG were recovered and repolymerized through esterification, pre-polycondensation and final polycondensation to generate high-quality PET (**Fig. 4a**, middle and right panels). The regenerated PET was subsequently processed by injection molding to produce solid PET specimens with crystallinities ranging from 26.7% to 39.1% (**Fig. 4a**, middle and right panels, **Supplementary Fig. 10**, **Extended Data Table. 2**). Upon a second cycle of enzymatic depolymerization, VenusPETase again efficiently degraded the recycled PET and generated high levels of PET-derived monomers (**Fig. 4a**, right panel), demonstrating that the recycled material remained fully compatible with enzymatic depolymerization and validating the feasibility of repeated enzymatic closed-loop recycling.

**Fig. 4.**
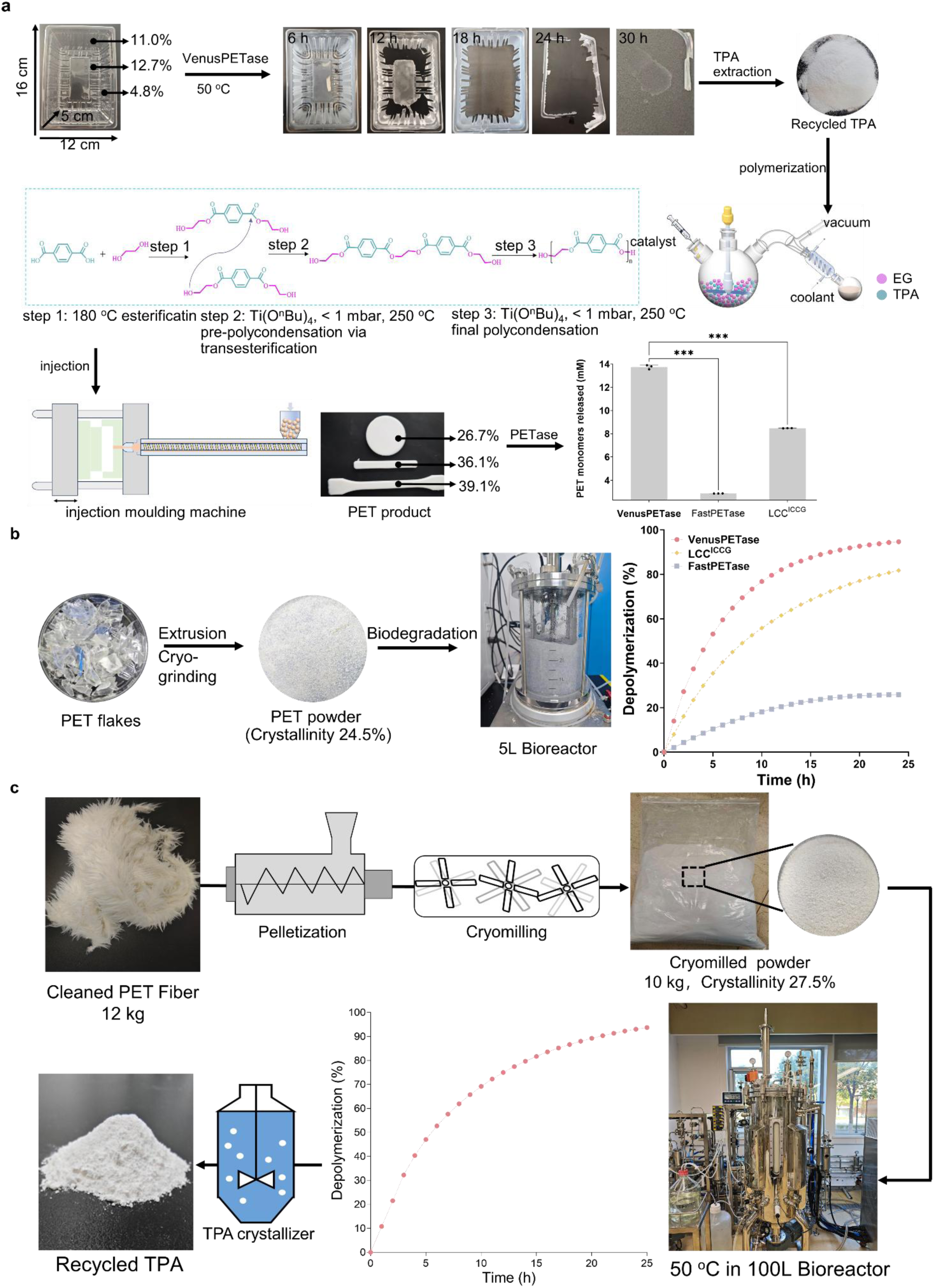
VenusPETase enables efficient enzymatic PET depolymerization and closed-loop recycling in scalable systems. **a**, Time-course images showing enzymatic degradation of a molded PET plastic block (12 cm × 16 cm × 0.5 cm) with crystallinity gradients (4.8%–12.7%) by VenusPETase at 50 °C. Without mechanical stirring, the PET block was degraded to near completion within 30 h, leaving only trace solid remnants.Middle panel illustrates a three-step chemical repolymerization process using enzymatically recovered monomers: (1) esterification of terephthalic acid (TPA) and ethylene glycol (EG) at 180 °C, (2) Ti(OBu)_4_-catalyzed transesterification under vacuum (<1 mbar, 250 ℃), and (3) final polycondensation to form poly(ethylene terephthalate). The resulting PET polymer was processed into shaped products via injection molding. As shown in the right panel, under standard reaction conditions (3 mg_enzyme_g_PET_⁻¹, 100 mM phosphate buffer, pH 8.0), VenusPETase-mediated PET degradation exhibited significantly higher monomer release (quantified by UPLC) compared to both FastPETase and LCC^ICCG^. Notably, VenusPETase and FastPETase reactions were conducted at their optimal temperature of 50°C, while LCC^ICCG^ was tested at its optimal temperature of 65°C. Bar heights represent mean ± s.d. of three biological replicates. **b**, Comparison of depolymerization performance in a 5 L bioreactor. Real recycled plastic bottle flakes were pretreated and processed into powders with a crystallinity of 24.5%. Depolymerization was performed in a 5 L bioreactor using VenusPETase, FastPETase, and LCC ^ICCG^ at 20% substrate loading and 1.5 mg_enzyme_ g_PET_⁻¹ for 24 h. Reactions with VenusPETase and FastPETase were conducted at 50 °C, whereas reactions with LCC^ICCG^ were conducted at 65 °C. PET depolymerization percentages were calculated from NaOH consumption. **c**, Demonstration of the potential industrial application of VenusPETase. Waste textiles were pretreated and converted into powder substrates with a crystallinity of 27.5%. Enzymatic depolymerization was performed in a 100 L bioreactor at 20% substrate loading and 1.5 mg_enzyme_ g_PET_⁻¹, using tap water as the reaction medium, for 25 h. PET depolymerization percentages were calculated from NaOH consumption, with the reaction pH maintained at 9.0.

A major challenge for industrial PET biorecycling is that post-consumer PET typically exhibits crystallinities of 30–40%, rendering it intrinsically resistant to enzymatic hydrolysis(*20, 37, 67*). Consequently, most previous PETase-based recycling processes have relied on extensive pretreatment to reduce crystallinity to around 10%(*33, 35, 37, 49, 68*), increasing both energy consumption and processing costs(*50, 69, 70*). Because VenusPETase preferentially degrades crystalline PET, we reasoned that efficient depolymerization could be achieved while preserving relatively high substrate crystallinity. To test this hypothesis, post-consumer PET bottle flakes were processed to increase surface area while retaining a crystallinity of 24.5% (**Supplementary Fig. 10**, **Extended Data Table 2**). VenusPETase was then benchmarked against LCC^ICCG^ and FastPETase in a 5 L bioreactor at 200g kg^-1^ substrate loading and an enzyme dosage of 1.5 mg _enzyme_ g_PET_^−1^, as the high-solids condition provides a more stringent test of catalytic performance while balancing operational cost and degradation efficiency(*60, 71*) (**Fig. 4b**). Depolymerization progress was monitored by NaOH consumption during pH-controlled hydrolysis(*33, 35, 49*). VenusPETase achieved >90% PET conversion within 17 h and near-complete depolymerization within 24 h, substantially outperforming LCC^ICCG^ and FastPETase which reached only 72% and 24% conversion at 17 h, respectively (**Extended Data Fig. 9**). Given that techno-economic analyses from the National Renewable Energy Laboratory predict that increasing PET conversion from 90% to 99% can reduce the minimum selling price of TPA by approximately 10% (US$0.2/kg)(*60*), the high conversion efficiency of VenusPETase can substantially improve the economic viability of enzymatic PET recycling.

To further assess industrial scalability, we extended process validation to a pilot-scale bioreactor following recently proposed standardization frameworks for PETase evaluation(*21*). Because polyester textiles constitute a major and rapidly growing PET waste stream(*2, 72, 73*), we selected post-consumer textile waste as a representative feedstock. Kilogram-scale textile waste was mechanically pretreated while maintaining a crystallinity of 27.5% (**Supplementary Fig. 10**, **Extended Data Table 2**) and subjected to enzymatic depolymerization in a 100 L bioreactor using the same substrate loading and enzyme dosage employed in the 5 L system. VenusPETase achieved more than 90% depolymerization within 20 h (**Fig. 4c**). Downstream purification yielded recycled TPA with a purity exceeding 99.5% and an overall recovery yield of approximately 94% (**Supplementary Fig. 11**). These results demonstrate that VenusPETase enables efficient, high-conversion depolymerization of high-crystallinity post-consumer PET at pilot scale and supports its potential application in industrial closed-loop recycling processes.

## Discussion

VenusPETase establishes a robust and scalable platform for enzymatic depolymerization of crystalline PET. Although substantial progress has been made in PETase engineering(*26-28, 30, 37, 40, 49, 66*), efficient degradation of highly crystalline PET remains a central challenge for enzymatic recycling because the densely packed polymer architecture restricts enzyme accessibility to ester bonds and substantially reduces hydrolysis efficiency. Consequently, many engineered PETases exhibit reduced catalytic activity on crystalline substrates, incomplete depolymerization, accumulation of intermediates such as mono(2-hydroxyethyl) terephthalate (MHET), or insufficient operational stability under industrially relevant conditions(*20, 22, 23, 33, 62, 74*). VenusPETase simultaneously addresses several of these long-standing limitations by combining high catalytic efficiency, broad crystallinity tolerance, exceptional thermostability, and near-complete conversion of PET hydrolysis products into TPA. VenusPETase maintains efficient depolymerization across PET substrates spanning a broad crystallinity range (8%–50%), including highly crystalline and post-consumer materials that are typically resistant to enzymatic hydrolysis(*20, 62*). In particular, the high TPA fraction substantially reduces intermediate accumulation and may simplify downstream purification and repolymerization processes, while the ability to process highly crystalline PET could reduce the need for crystallinity-based waste sorting or aggressive thermomechanical pretreatment(*31, 50*). Importantly, VenusPETase retains high catalytic performance under moderate operational temperatures and demonstrates efficient depolymerization at pilot and industrially relevant scales, including complete depolymerization of post-consumer crystalline PET in a 100 L bioreactor. Together, these results substantially extend the scope of enzymatic PET recycling beyond low-crystallinity laboratory substrates(*33, 35, 37, 49*) toward industrially relevant plastic waste streams and establish VenusPETase as a promising platform for scalable closed-loop PET recycling.

Mechanistically, the enhanced catalytic performance of VenusPETase arises from coordinated optimization across multiple structural and dynamic scales. Structural and MD analyses suggest that increased electrostatic surface potential collectively promote productive engagement with PET substrates, particularly highly crystalline PET. At the catalytic center, VenusPETase maintains substrate conformations that favor transition-state stabilization and lowers the hydrolytic energy barrier during both acylation and deacylation steps. Unlike many previously engineered PETases that rely primarily on disulfide-bond introduction to improve thermostability(*28, 35, 37, 38, 49*), VenusPETase achieves enhanced performance through dynamic modulation and active-site microenvironment remodeling while preserving the overall α/β-hydrolase architecture. These findings suggest that simultaneous optimization of conformational dynamics, substrate accessibility, and catalytic geometry may represent an effective strategy for overcoming the persistent trade-off between catalytic efficiency and thermostability in polyester hydrolases.

Beyond the performance of VenusPETase itself, this work establishes PET-Flow as a data-efficient framework for AI-guided PETase engineering. Current machine learning approaches for enzyme design frequently face major challenges in exploring combinatorial sequence space while jointly optimizing multiple functional properties, particularly when only limited experimental data are available(*26, 35, 38, 49*). PET-Flow addresses this problem through a two-round design strategy that integrates zero-shot mutant-effect prediction with supervised combinatorial optimization under thermostability constraints. In the first round, a protein language model fine-tuned on PETase homologs enabled efficient identification of beneficial single-site substitutions without requiring prior experimental mutant data. In the second round, experimentally validated single-site mutants were used to train supervised models that explored higher-order mutational combinations while filtering unstable variants. This strategy substantially reduced the experimental search space while enabling progressive optimization of catalytic activity, thermostability, and product selectivity within a single scaffold. PET-Flow also is capable of partially overcoming epistatic barriers that frequently limit conventional directed evolution approaches. More broadly, these results demonstrate how protein language models and limited-data learning strategies can be integrated to navigate complex sequence-structure-function landscapes for enzyme engineering.

More broadly, this work highlights the emerging potential of AI-guided PETase engineering for addressing major challenges in polymer recycling. The ability to computationally optimize catalytic efficiency, thermostability, substrate scope, and product selectivity within a unified design framework may accelerate the development of next-generation biocatalysts for synthetic polymer depolymerization. Beyond PET, similar strategies may be applicable to other recalcitrant polyesters and industrially relevant polymers.

## Method

### ProtSSN Fine-tuning and Zero-shot Scoring

#### Homologous sequence collection

To fine-tune ProtSSN for PETase-specific mutant effect prediction, we first curated a dataset of PETase homologous sequences. We used the all known PETase sequence as the query and performed a sequence similarity search against the UniProtKB database using Jackhmmer(*75*) with an E-value threshold of 0.00001. After removing sequences with length outside the range of 10 to 1024 amino acids and filtering redundant sequences at 30% identity, we obtained 179,753 homologous sequences. We predicted their structures using ESMFold(*76*), resulting in 179,753 sequence-structure pairs for fine-tuning.

#### Fine-tuning procedure

ProtSSN utilizes both protein sequence and structure as input. We fine-tuned ProtSSN using a masked language modeling objective on the collected sequence-structure pairs. The model was trained to predict randomly masked residues (masking rate: 15%) based on their structural and sequential context. Fine-tuning was performed for 5 epochs with a learning rate of 0.0001, global batch size of 256, and the AdamW optimizer. The fine-tuned model is referred to as ProtSSN-evotuned.

#### Zero-shot fitness scoring

For mutant fitness prediction on KbPETase, we used the experimentally determined crystal structure (PDB: 9IW9) of the wild-type. For each candidate single-site mutant, we computed a fitness score using ProtSSN-evotuned based on the ratio of predicted amino acid probabilities. Specifically, given a wild-type sequence ***S*** with structure ***G*** and a mutation at position ***i*** from wild-type residue ***a*** to mutant residue ***b***, the fitness score was calculated as:

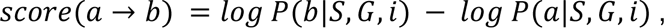

where *P*(· |*S*, *G*, *i*) represents the predicted probability of an amino acid at position *i* given the sequence and structural context. A score greater than 0 indicates that the mutant residue is predicted to be more favorable than the wild-type residue at that position, suggesting the mutation is more consistent with evolutionary constraints learned from homologous sequences. Conversely, a score less than 0 suggests the mutation is evolutionarily disfavored. All 4,826 possible single-site mutants (19 substitutions × 254 mutable positions) were scored and ranked in descending order by their fitness scores.

### Ankh + MLP Supervised Learning Model

#### Sequence encoding

Mutant sequences were encoded using Ankh(*55*), a protein language model pretrained with auto-regressive masked language modeling on UniRef50(*77*), For each mutant sequence, we extracted the final-layer hidden representations and computed the mean pooling across all residue positions to obtain a fixed-dimensional embedding vector of size 768.

#### Model architecture

The supervised prediction model consisted of the frozen Ankh encoder followed by a multi-layer perceptron (MLP) head. The MLP architecture comprised one fully connected layer with hidden dimensions of 100, ReLU activation functions. Two separate models were trained: one for predicting catalytic activity and one for predicting melting temperature (T_m_).

#### Training procedure

Models were trained on the Round 1 experimental data comprising 20 single-site mutants with measured activity and T_m_ values. The dataset was split into a training set (15 mutants) and a validation set (5 mutants); the specific mutant assignments for each split are provided in **Supplementary Table 3**. We used mean squared error (MSE) as the loss function and trained for a maximum of 50 epochs with early stopping based on validation loss (patience = 5 epochs). The Adam optimizer was used with an initial learning rate of 1 × 10^-4^ and a cosine annealing learning rate schedule.

#### Inference

For multi-site mutant design, we used the trained models to predict activity and Tm for all candidate variants generated during the stepwise combinatorial search. Candidates were filtered by a thermostability threshold (predicted T_m_ ≥ 80 °C), and the remaining variants were ranked by predicted activity.

### PET-Gym Benchmark Construction and Evaluation

To systematically evaluate zero-shot predictors and supervised learning methods for PETase mutant effect prediction, we constructed PET-Gym, a comprehensive benchmark comprising experimental data from multiple PETase variants.

#### Data collection

PET-Gym aggregates published mutational data from five PETase enzymes: BhrPETase(*49*), CaPETase(*35*), IsPETase(*26*), KubuPETase(*38*), and MipaPETase(*38*). For each PETase, we collected single-site and multi-site mutant data with measured catalytic activity and/or melting temperature (T_m_). The complete dataset comprises 290 single-site mutants and 258 multi-site mutants across the five enzymes (detailed statistical information **Supplementary Table 4** and data are in the code repository).

#### Benchmark tasks

We defined two evaluation settings to assess different prediction manners: (i) Zero-shot evaluation (PET-Gym-zero-shot). This task evaluates the ability of unsupervised or zero-shot methods to predict mutant effects without any mutant training data. For each PETase in the benchmark, we computed the Spearman correlation coefficient between predicted fitness scores and experimentally measured activity or Tm across all single-site mutants. We evaluated several zero-shot methods, including ProtSSN(*47*), MIF-ST(*78*), ESM-1v(*79*), S3F(*80*), VenusREM(*81*), ESM-2(*76*)and SaProt(*82*). Results are shown in **Extneded data Fig 3**. (ii) Supervised evaluation with generalization to multi-site mutants (PET-Gym-single-to-high). This task assesses whether supervised models trained exclusively on single-site mutant data can generalize to predict the effects of multi-site (higher-order) mutants. For each PETase, single-site mutants were used as the training set, and multi-site mutants (double, triple, and higher-order) served as the held-out test set. We evaluated combinations of protein language model encoders (ESM2(*76*), Ankh(*55*), xTrimoPGLM(*83*)) and machine learning algorithms (**Supplementary Table 5**). Model performance was measured by Spearman correlation and root mean squared error (RMSE) on the test set. Results are presented in **Extneded data Fig 3**.

#### Evaluation results

In the zero-shot setting, ProtSSN achieved the highest average Spearman correlation (Spearman’s ρ = 0.374) across the five PETases, outperforming ESM-1v (Spearman’s ρ = 0.258) and ESM-IF1 (Spearman’s ρ = 0.329). Fine-tuning on homologous sequences (ProtSSN-evotuned) further improved performance to 0.435 (Spearman’s ρ), demonstrating the benefit of domain-specific adaptation. In the supervised single-to-multi-site setting, the combination of Ankh embeddings with an MLP head yielded the best generalization performance (Spearman’s ρ = 0.596), outperforming other models. These results guided our selection of ProtSSN-evotuned for Round 1 zero-shot scoring and Ankh+MLP for Round 2 supervised prediction in the PET-Flow pipeline.

### PET-Flow for evolve KbPETase

#### Overview

PET-Flow is a two-round deep-learning guided pipeline for PETase engineering that integrates zero-shot mutation effect prediction with supervised learning-based combinatorial optimization. Round 1 employs zero-shot prediction to screen single-site mutations, while Round 2 leverages supervised models trained on Round 1 data to systematically explore the multi-site combinatorial space.

#### Single-site mutant select strategy

From the zero-shot fitness scores generated by ProtSSN-evotuned, we implemented a position-diversity constraint to ensure comprehensive coverage across the protein sequence for downstream combinatorial design. Specifically, we ranked all single-site mutants by their predicted fitness scores and iteratively selected the highest-scoring mutant at each position, retaining at most one mutation per residue. This procedure yielded a set of 20 single-site mutants for experimental validation. The predicted fitness scores and experimentally measured values (activity and T_m_) for these 20 mutants are presented in **Supplementary Table 1**.

#### Multi-site mutant select strategy

The n-site mutant selection procedure is formalized as follows (selecting one n-site mutant per iteration):

i. Candidate generation: Starting from the selected (n−1)-site mutant, we enumerated all remaining single-site mutations and combined each with the (n−1)-site backbone to generate n-site candidates. This yielded (20 − n + 1) combinations per iteration. Note: For n=2, we initialized the search from mutant R201L, which exhibited the highest performance among single-site variants.
ii. Property prediction: For each candidate variant, we predicted catalytic activity and Tm using the trained Ankh+MLP models.
iii. Stability filtering: We retained only candidates whose predicted T_m_ exceeded 80 °C.
iv. Best mutant selection: Among filtered candidates, we selected the mutant with the highest predicted activity.

This iterative procedure was executed from two-site through fifteen-site designs. When no candidate at a given iteration satisfied the stability threshold, that mutation order was skipped and the search continued to the next iteration using the same backbone variant. This occurred at the eight-site and nine-site steps, ultimately yielding 12 multi-site variants (rather than 14) for experimental validation. The selected multi-site mutants and experimentally measured values (activity and Tm) for these mutants are presented in **Supplementary Table 2**.

#### Stability Threshold Rationale

The stability threshold was set at T_m_ > 80 °C based on the operational temperature range of industrial PET hydrolysis (65–72 °C) plus a thermal stability margin. The maximum mutation order was capped at 15. At iterations 8 and 9, no candidates met the stability criterion; accordingly, the algorithm bypassed these steps and proceeded to the next iteration, resulting in 12 multi-site variants for experimental characterization.

### PET membrane model construction for molecular dynamics simulations

For the low-crystallinity PET model (APET), we first optimized the PET monomer structure by Gaussian 16 Rev. C.01(*84*) at B3LYP/def2SVP(*85, 86*) level with D3(BJ) empirical dispersion(*87*). Then the partial charges were set to fit the electrostatic potential generated with B3LYP/def2TZVP by RESP2(0.5) method(*88*) through Multiwfn(*89*) integrated with LIBRETA(*90*). The MD simulations were performed with Gromacs 2025.2(*91*) using the general Amber force field (GAFF)(*92*). The modelling protocol was presented in **Supplementary Figure 4**. The PET chain contains 100 repeated MHET units. Initially, 14 PET chains were placed in a 10 × 10 × 250 nm large periodic cubic box and simulated for 50 ns at 800 K using V-rescale thermostat(*93*) at NVT ensemble. Then the system underwent NVT simulations with two walls at z = 0 and z = box boundaries, composed of sp^2^ C pure aromatic atoms (ca atom type in GAFF) with 12-6 wall-type. The system was deformed with 0.1 nm/ps speed along the Z-axis to compress the PET complex into a 10 × 10 × 3.5 nm membrane. To equilibrate the different PET chains, we extended the simulations for 100 ns at 800 K and subsequent 100 ns simulation at 323 K. Finally, the PET membrane was solvated in a 10 × 10 × 15 nm water box, with TIP3P(*94*) water molecules, and equilibrated for 100 ns at 323 K and 1 bar (NPT ensemble), with V-rescale thermostat and C-rescale(*95*) barostat and smooth PME (SPME) electrostatic method(*96*). The finally equilibrated PET structure was taken as APET model in the subsequent simulations.

For the high-crystallinity PET model (CPET), as presented in **Supplementary Figure 4**, we first optimized the cell of PET crystal by CP2K(*97*), and then expanded the cell to 10 × 10 × 5 nm. The initial crystal cell was constructed according to ref(*98*). The optimization was carried out with PBE method(*99*) and DZVP-MOLOPT-SR-GTH basis set(*100*), with k-point set as “4 3 2” at x, y, and z directions, separately. The input file for CP2K was generated by Multiwfn(*101*). During subsequent MD simulation, the PET molecule was treated as a periodic molecule, and the topology was built by Sobtop 1.0(dev 5)(*102*) with 1.2*CM5 atom charge obtained through Multiwfn(*89*). The system was first equilibrated for 50 ns in NVT ensemble, and then solvated and underwent 50 ns MD simulations in NPT ensemble. Other simulation conditions were the same as APET construction. The finally equilibrated PET structure was taken as CPET model in the subsequent simulations.

After constructing the two models, we verified the structural validity by investigating the key dihedrals in the PET chains (**Supplementary Figure 5**). The dihedrals were calculated by the Matplotlib python package. Visualization of the PET models were accomplished using PyMOL(*103*) and VMD(*104*).

### Molecular dynamics simulations for KbPETase and VenusPETase

The crystal structures of KbPETase (PDB ID: 9IW9) and VenusPETase (PDB ID: 9VJ7) were used for MD simulations. After protonation states decision of titratable residues and removing all crystallographic waters, we performed MD simulations of both enzymes with substrates by Gromacs 2025.2(*91*) with Amber ff14SB force-field(*105*) in a water box containing APET/CPET membranes. Both enzymes were placed above the membrane with a 1 nm distance. Na^+^ and Cl^−^ ions were added to neutralize the protein charge at 0.15 M concentration. The system underwent energy minimization using the steepest descent method for 2000 steps. Subsequently, the system was equilibrated for 10 ns at 323.15 K and 1 bar under the NPT ensemble. Then each system (APET/*Kb*PETase, APET/VenusPETase, CPET/*Kb*PETase, CPET/VenusPETase) underwent 100 ns production simulations. The cutoff radius for neighbor list searching and nonbonded interactions was taken to be 1.0 nm, and all bonds were constrained using the LINCS algorithm(*106*). Visualization of enzyme structures was accomplished using PyMOL(*103*) and VMD(*104*). Curvature and dihedral analysis were carried out by the Membrane Curvature tool utilizing MDAnalysis 2.9.0 python package(*107*). Besides APET/CPET substrates, we also built 2PET model substrate (two linked MHET molecules). The topology of 2PET was built by Sobtop 1.0(dev 5)(*102*) with RESP2(0.5) charge(*88*). The 2PET molecule was docked to KbPETase and VenusPETase at S128, H206 and D176 catalytic triad by DSDP(*108*) and each binding pose with the highest affinity was selected. After 2000 steps of energy minimization, each system underwent 50 ns simulation at 323.15 K and 1 bar. Each simulation was repeated for three replicas to ensure statistical significance. The binding energy was analyzed by s_mmpbsa program 0.10.3(*109*). Furthermore, metadynamics is performed by integrated COLVARS module(*110*), based on the representative structure of previous MD simulations, and the collective variables (CV) were set as S128^Oγ^ − PET ^ester−C^ distance (d_1_) and S128^Oγ^ − H206^Nδ^ distance (d_2_). The hill width was set to 0.25 nm, with hill weight as 0.01 kJ/mol. Each system underwent 10 ns metadynamics simulation, with COLVARS trajectory frequency as per 100 steps.

### First-Principles Molecular dynamics simulations for KbPETase and VenusPETase

Based on the equilibrated KbPETase/2PET and VenusPETase/2PET complex structures, we extracted the active site residues (S128, D176, H206, Y58, and M129) together with 2PET substrate and capped with hydrogen atoms. Then we performed FPMD simulations using CP2K 2024.1 software(*97*). The simulations employed GFN1-xTB(*111*) method at 345.15 K with a time step of 0.2 fs, with input files generated by Multiwfn(*89*). During simulations, all Cα atoms were fixed. Two distances were set as CVs: distance between the Oγ atom of S128 and the carbonyl carbon of 2PET (Oγ^S128^–C^ester^, *d*_1_), and the distance between the carbonyl carbon of 2PET and ester/water oxygen (C^ester^–O^ester/water^, *d*_2_). The system first underwent 2000 steps of equilibration without bias, and the CV standard deviation was calculated and set as the bias potentials width in the following stage of 10000 steps production phase, with a gaussian hill height of 1.5 kcal/mol, under self-consistent continuum solvation condition. Energy values of different reaction states were subsequently estimated using the kernel density estimation method.

### Plasmid construction

Codon-optimized coding sequences of KbPETase and its mutants (**Supplementary Table 6**) predicted by PET-Flow were synthesized by Sangon Biotech (Shanghai, China) and subcloned into the pET-28a (+) expression vector via *Nde*I and *Xho*I restric-tion sites, incorporating an N-terminal hexahistidine (His_6_) tag. TurboPETase(*49*), LCC^ICCG^(*37*), PES-H1^L92F/Q94Y^(*39*), and FastPETase(*2*) were individually inserted into the pET-28a(+) *Nde*I and *Xho*I digestion, also retaining an N-terminal His_6_ tag. DepoPETase(*27*), ThermoPETase(*40*), CaPETase^M9^(*35*), Kubu-P^M12^(*38*), and HotPETase(*28*) were cloned into the pET-21a (+) using *Nde*I and *Xho*I restriction sites, followed by fusion to a C-terminal His_6_ tag.

### Recombinant protein expression and purification in *E. coli*

The recombinant plasmids were transformed into *E. coli* Rosetta-gami B (DE3) competent cells (Weidibio, China). The transformed cells were grown in lysogeny broth (LB) medium supplemented with 100 g/mL Kanamycin at 37 ℃ with shaking at 220 rpm until OD_600_ reached 0.6–1.0, then protein expression was induced by adding 0.6 mM isopropyl β-D-1-thiogalactopyranoside (IPTG) and continued at 16 ℃ for 18–20 h (220 rpm). Cells were harvested by centrifugation at 3800 × g for 25 min at 4 ℃. The cell pellets were resuspended in a lysis buffer (25 mM Tris-HCl, 500 mM NaCl, pH 7.4) and disrupted by ultrasonication (Scientz, China). Lysates were clarified by centrifugation at 17,400 × g for 30 min at 4 ℃, then the clarified supernatants were applied to a Ni-NTA agarose column (Smart Lifesciences, China, Ni NTA Beads 6FF: SA005500) pre-equilibrated with lysis buffer. After washing, bound proteins were eluted with an elution buffer (25 mM Tris-HCl, 500 mM NaCl, 250 mM imidazole, pH 7.4). All purification steps were performed at 4 ℃ to maintain protein stability. Purified proteins were analyzed by 15% SDS-PAGE and visualized with Coomassie Brilliant Blue R-250 staining. The enzymes were concentrated using Amicon Ultra centrifugal filter units (10 kDa MWCO, Millipore, US) and stored at −80 ℃ in 15% glycerol for long-term preservation.

### Differential scanning fluorimetry (DSF)

Protein melting temperature (T_m_) was determined using the DSF method with the Protein Thermal Shift Dye Kit (Thermo Fisher, US). To prepare the reaction mixture, 1.0 µL of SYPRO Orange Dye (SUPELCO, US) was diluted in 49 µL of lysis buffer (25 mM Tris-HCl, 500 mM NaCl, pH 7.4)(*28, 36, 112*). Next, 1 µL of the diluted dye was combined with 19 µL of protein solution at a protein concentration of 0.1 mg/mL. DSF experiments were conducted using the LightCycler 480 Instrument II (Roche, US). The reaction mixture was equilibrated at 25 °C and then heated to 99 ℃ at a rate of 0.05 ℃/s, with a 2 minutes hold at the final temperature.

### Enzymatic assay of KbPETase and its mutants

The PET-degrading activity of KbPETase and its mutants was quantitatively evaluated using crystalline PET powder (Cry-PET; Goodfellow Cambridge Ltd, Cat. No. ES306000, 39.8% Crystallinity)(*35*). The crystalline PET powder was used for assay without any pretreatment. Standard enzymatic reactions were conducted in 1 mL of 50 mM glycine-NaOH buffer (pH 9.0) containing 500 nM enzyme and 15 mg Cry-PET powder at 50 ℃ for 24 h. The reactions were terminated by heating the mixture at 95 ℃ for 10 min(*36, 113*). Each sample was diluted to fall within the linear detection range for TPA and MHET. After filtration through a 0.22 μm filter, the assay solution was analyzed by ultra-performance liquid chromatography (UPLC)(*28, 36*).

### Comparing VenusPETase with other PETases under their respective optimal buffer

The PET-degrading assay was performed by incubating 15 mg of Cry-PET powder with 1 mL of reaction buffer at 50, 60 and 65 ℃ for 24 h. Depolymerization was initiated by adding purified PETase to a final loading of 500 nM. For LCC^ICCG^ (*37*), TurboPETase(*49*), FastPETase(*2*), KubuP^M12^(*38*) and PES-H1^L92F/Q94Y^(*39*), reactions were carried out in 100 mM potassium phosphate buffer (pH 8.0). For ThermoPETase(*40*) and CaPETase^M9^(*35*), reactions were performed in 50 mM glycine−OH buffer (pH 9.0). For VenusPETase and HotPETase (*28*), reactions were performed in 50 mM glycine−OH buffer (pH 9.2). For DepoPETase(*27*) reactions were assayed in 100 mM glycine−OH buffer (pH 9.0). The reactions were terminated by heating the mixture at 95 ℃ for 10 min. Each reaction was diluted to fall within the linear detection range for TPA and MHET. After filtration through a 0.22 μm filter, the assay solution was analyzed by UPLC (*28, 36*).

### Evaluation of degradation efficiency on PET with different crystallinity

PET samples with different degrees of crystallinity were used as substrates. Amorphous PET films were obtained from a commercial source (Goodfellow, Cat. No. ES301445, England) (*2, 36*). Low- to medium-crystallinity PET samples were prepared from post-consumer PET waste, including low-crystallinity PET from disposable fruit containers and medium-crystallinity PET from the sidewalls of water bottles. High- crystallinity PET substrates were derived from colored PET flakes collected during recycling processes. All PET materials were cryogenically ground to obtain particles with diameters smaller than 200 µm. The crystallinity of each substrate was determined prior to enzymatic assays and subsequently used as the basis for comparative analysis. Enzymatic depolymerization reactions were performed by incubating 15 mg of PET powder in 1 mL of buffer containing 100 µL of enzyme solution (stock concentration: 0.5 mg/mL) for 72 h. Reactions involving LCC^ICCG^ and FastPETase were carried out in 100 mM potassium phosphate buffer (pH 8.0) at 65 ℃, whereas reactions with VenusPETase were conducted in 100 mM glycine–NaOH buffer (pH 9.0) at 50 ℃. The reactions were terminated by heating the mixture at 95 ℃ for 10 min. Reaction mixtures were diluted to ensure that TPA and MHET concentrations were within the linear detection range, filtered through 0.22 µm membrane, and analyzed by UPLC(*28, 36*).

### Depolymerization of post-consumer PET plastics by VenusPETase

A large, untreated, transparent pc-PET container (fruit packing type, approximately 10.4 g) was treated by 500 nM of VenusPETase in 100 mM of Glycine-NaOH buffer (pH 9.2) at 50 ℃(*2*). The whole piece of transparent PET container was fully submerged in 2.5 L of enzyme solution and completely degraded after 32 h.

### X-ray diffraction analyses for PET crystallinity

The crystallinity of PET samples was characterized by XRD using a Bruker D8 ADVANCE diffractometer (Bruker, Germany) and a PANalytical Empyrean system. Measurements were per- formed with Cu Kα radiation (λ = 1.5406 Å) generated at 40 kV and 40 mA, employing a nickel filter and a PIXcel3D detector. Diffraction patterns were collected in continuous-scan mode over a 2θ range of 10-80° with a scan rate of 0.042°/s and a 0.5° divergence slit. All measurements were conducted at ambient temperature.

### Enzyme heat-inactivation assay(*35*)

Thermal stability assays for PET degradation were conducted to assess the heat inactivation profiles of VenusPETase, FastPETase, and LCC^ICCG^. Each enzyme (500 nM) was prepared in its respective optimal buffer—50 mM glycine-NaOH (pH 9.0) for VenusPETase, and 100 mM potassium phosphate (pH 8.0) for FastPETase(*2*) and LCC^ICCG^(*37*). The enzyme solutions were incubated at 50 °C and 65 °C for 72 h. Subsequently, 15 mg of Cry-PET powder(*35*) was added to each preincubated enzyme solution, and the reaction mixtures were incubated at their respective optimal temperatures (50 °C for VenusPETase and FastPETase, 65 °C for LCC^ICCG^) for an additional 24 h. The reactions were terminated by heating the mixture at 95 °C for 10 min. The resulting hydrolysates were filtered through a 0.22 µm PVDF syringe filter and analyzed by UPLC.

### Pretreatment of post-consumer PET

Post-consumer PET was collected from a municipal recycling facility in Shanghai, China and processed into small flakes by a commercial waste handler. The samples were subsequently amorphized using a twin- screw extruder (Xinshuo, Shanghai, China) operated at 265 °C in the extrusion zones, 285 °C in the melt pump, and 285 °C in the screen changer zones. The resulting PET strands were further micronized at room temperature using a JXYD-2000 grinder (Jingxin, Shanghai, China). After sieving, PET powders with particle sizes below 400 µm were obtained.

### Comparing the depolymerization performance of VenusPETase and other PETases at high PET loading

The large-scale depolymerization capacity of VenusPETase was evaluated using a 5 L bioreactor (Baotech, Shanghai, China) containing with 330 g of mechanically pretreated PET powder (particle size 400 µm, average crystallinity of 24.5%) suspended in 1.6 L of 100 mM potassium phosphate buffer (pH 8.0) [2, 31, 36]. The reaction mixture was supplemented with 49.5 mL of enzyme solution comprising 495 mg of purified PETases (10 mg/mL in 20 mM Tris-HCl buffer, 500 mM NaCl, pH 7.4). The system was maintained at 50 ± 0.5 °C for FastPETase(*2*) and VenusPETase or 65 ± 0.5 °C for LCC^ICCG^ (*37*) with continuous agitation at 250 rpm, while the pH was automatically stabilized at 8.05 ± 0.1 through titration with 6 M NaOH. Real-time monitoring of depolymerization was performed by recording NaOH consumption, which directly correlates with TPA release.

### Bioreactor-scale evaluation of VenusPETase for PET depolymerization

To assess the industrial-scale applicability of VenusPETase, PET depolymerization was performed in a 100-L bioreactor (Bailun, Shanghai, China). Pretreated post-consumer PET powder (particle size 400 µm, average crystallinity of 27.5%, 10 kg) was added to the reactor together with 48.5 L of tap water, and the mixture was heated to 50–60 °C. VenusPETase (10 mg/mL in 20 mM Tris-HCl buffer, 500 mM NaCl, pH 7.4) was then introduced at a total amount of 15 g. The final reaction conditions corresponded to a substrate loading of approximately 20% (w/v) and an enzyme dosage of approximately 1.5 mg g_PET_^−1^. During the reaction, the temperature was maintained at 50.0 ± 0.5 °C, the agitation speed was set to 250 rpm, and the pH was automatically maintained at 8.05 ± 0.1 using 6 M NaOH. The progress of PET depolymerization was continuously tracked by measuring NaOH consumption, which directly reflected the release of TPA.

### Chromatographic analysis of PETase reactions

The quantification of PET degradation products was performed using an Agilent 1290 Infinity II UPLC system (Agilent, USA) equipped with a UV detector set at 260 nm. Separation was achieved on a Kinetex XB-C18 column (50 × 2.1 mm, 5 µm, 100 Å; Phenomenex, USA) under a stepped-isocratic gradient. The mobile phases consisted of 0.1% formic acid in water and acetonitrile, delivered at a flow rate of 1.1 mL/min. Samples (3 µL injection volume) were first eluted with 13% B for 52 s to resolve MHET and TPA, followed by a 33 s gradient to 95% B to separate higher oligomers. The total run time was 2 min, including column re-equilibration. Analytes were identified by retention times with authentic standards (TPA: ∼0.4 min; MHET: ∼0.6 min; BHET/oligomers: ∼1.0–1.2 min), and quantification was performed via external calibration curve (*28, 36*).

### Michaelis-Menten kinetics analysis of PETase

For kinetics analysis, enzymatic reactions using 4-nitrophenyl butyrate (*p*NPB) as the substrate were conducted in 96-well plates with a total volume of 100 µL(*36, 49*). The final *p*NPB concentrations ranged from 0.2 to 1.8 mM, while the enzyme concentration was maintained at 0.5 µg/mL. Each reaction mixture contained 10 µL of *p*NPB solution (dissolved in anhydrous ethanol), 80 µL of 10 mM potassium phosphate buffer (pH 8.0), and 10 µL of enzyme solution, and was incubated at 50 ℃ for 3 min. Reaction termination and product detection were performed as described in the enzymatic activity screening section. One unit of enzymatic activity was defined as the amount of enzyme required to hydrolyse 1 µM of pNPB per minute. Kinetic parameters (K_m_ and V_max_) were determined by nonlinear regression fitting of the Michaelis-Menten equation using Prism (GraphPad), based on initial velocity data obtained from triplicate measurements at varying substrate concentrations.

### Expression and purification of VenusPETase for crystallization

Competent *E. coli* BL21(DE3) were transformed with VenusPETase expression plasmid, cultured at 37 ℃ until the optical density at 600 nm reached 0.6–1.0, and induced with 0.2 mM IPTG at 18 ℃ overnight. Cells were harvested by centrifugation at 5,000 g for 15 min at 4 ℃ and lysed in buffer A (20 mM Tris-HCl pH 7.5, 500 mM NaCl, 5% glycerol, and 1 mM TCEP) by sonication. The lysate was clarified by centrifugation at 40,000 g for 60 min at 4 ℃. The supernatant was loaded onto a Ni^2+^-NTA affinity column pre-equilibrated with buffer A. Unbound proteins were removed by washing with buffer A containing 20 mM imidazole, and VenusPETase was eluted using a lin- ear imidazole gradient (60–300 mM). To remove the N-terminal His_6_ tag, 500 U of thrombin was added to the pooled elution fractions, and the mixture was dialyzed overnight against buffer A. The cleaved protein was passed through the Ni^2+^-NTA column again to remove His_6_ tag and uncleaved protein. The tag-free VenusPETase was further purified by HiTrap Heparin HP affinity column, followed by size-exclusion chromatography in buffer containing 20 mM Tris-HCl pH 7.5, 150 mM NaCl, 5% glycerol, and 1 mM TCEP. SDS-PAGE analysis confirmed that the final VenusPETase preparation was ∼95% pure.

### X-ray crystallography of VenusPETase

VenusPETase (26.5 mg/mL in 20 mM Tris-HCl pH 7.5, 150 mM NaCl, 5% Glycerol, and 1 mM TCEP) was crystallized by the sitting-drop vapor diffusion method at 4 ℃, using 100 nL protein solution mixed with 150 nL of reservoir buffer. High-quality crystals grew within six weeks in reservoir buffer containing 0.2 M potassium bromide, 0.2 M potassium thiocyanate, 0.1 M sodium acetate (pH 5.0), 3% w/v γ-PGA (Na^+^ form, LM), and 10% w/v PEG 2000 MME. For data collection, crystals were soaked in a cryo-protectant solution consisting of the reservoir buffer supplemented with 20% v/v ethylene glycol, and then flash-frozen in liquid nitrogen. Diffraction data were collected at beamline BL10U2 of the Shanghai Synchrotron Radiation Facility (Shanghai, China) at a wavelength of 0.97915 Å and processed using XDS(*114*) and Aimless(*115*). The AlphaFold-predicted model of VenusPETase was used as the search template for molecular replacement with Phaser(*116*). The structure was refined through iterative cycles of manual model building in Coot(*117*) and refinement with Phenix.refine(*118*). Ramachandran statistics were evaluated using MolProbit(*119*)(**Supplementary Table 7**). All structural figures were generated in PyMOL(*103*).

### Post-degradation processing and terephthalic acid recovery

Following enzymatic PET degradation, the reaction mixture was filtered, and the filtrate was collected for subsequent regeneration. Acid precipitation was then performed by adjusting the pH to 2.0 using concentrated H_2_SO_4_ (98%), yielding crude TPA crystals. The precipitate was collected by filtration, washed thoroughly with deionized water, and dried under vacuum at 65 °C for 24 h to obtain 170.5 g of TPA (purity 95% by determined by UPLC).

### Recycled TPA-derived PET synthesis and injection molding

Pre-dried terephthalic acid (TPA, 166.13 g, 1 mol; vacuum-dried at 120 °C for 12 h) and ethylene glycol (EG, 63.93 g, 1.03 mol; dried over 4 Å molecular sieves) were loaded into a 500 mL three-necked round-bottom flask fitted with a mechanical stirrer, nitrogen inlet and distillation assembly. The reactor was purged with high-purity nitrogen three times to eliminate residual air. The mixture was stirred to form a homogeneous slurry, then heated to 180 °C in an oil bath and held at this temperature for 5 h to conduct the esterification reaction. Titanium (IV) butoxide (Ti(OBu)_4_, 0.34 g, 0.001 mol) was added when the rate of water distillation decreased substantially. Subsequently, the system pressure was gradually reduced to below 1 mbar to remove excess free ethylene glycol and the pre-polycondensation was carried out via transesterification at 250 °C for 2 h. The chain propagation stage was continued at 250 °C for 4 h under high vacuum to accomplish the final polycondensation. A notable rise in melt viscosity was observed during this stage. Upon completion of the reaction, nitrogen was admitted to restore atmospheric pressure. The molten product was quenched in deionized water, and the resultant solid was collected and dried at 80 °C for 24 h prior to further characterization (yield: 192.38 g).

The recycled terephthalic acid (TPA)-derived PET synthesized in this work was fabricated into standard molded specimens via conventional injection molding. A segmented barrel temperature strategy was adopted to ensure sufficient polymer melting and uniform melt fluidity, with the processing temperatures set as follows: 210 °C for the rear barrel zone, 240 °C for the middle barrel zone, 270 °C for the front barrel zone, and 260 °C for the nozzle. The injection process was performed at a constant injection pressure of 65 bar with a holding duration of 3 s to guarantee complete mold filling and compact specimen formation. After full demolding, the obtained injection-molded parts were naturally cooled to ambient temperature in atmospheric air to eliminate internal residual stress and stabilize the morphological structure of the polymer samples.

### Statistical analysis

Statistical analysis was performed in Prism (GraphPad). Statistical tests were one-sided or two-way ANOVA followed by Dunnett’s multiple comparisons test.

## Supplementary files

Supplementary information includes the raw data from the protein thermostability assay and the plasmid construct of VenusPETase. Supplementary file 1 includes the crystal structure of VenusPETase. Supplementary file 2 presents the videos of VenusPETase depolymerizing PET substrates.

## Acknowledgments

This work was supported by the National Key Research and Development Program of China (2024YFA0917603); Shanghai Municipal Science and Technology Major Project; the AI for Science Program, Shanghai Municipal Commission of Economy and Informatization (2025-GZL-RGZN-BTBX-02009); the Computational Biology Key Program of Shanghai Science and Technology Commission (23JS1400600); Shanghai Jiao Tong University Scientific and Technological Innovation Funds (21X010200843); Science and Technology Innovation Key R&D Program of Chongqing (CSTB2022TIAD-STX0017, CSTB2024TIAD-STX0032); the Student Innovation Center at Shanghai Jiao Tong University, and Shanghai Artificial Intelligence Laboratory. We thank Teacher Wanping Zhou from the Core Facility and Technical Service Center for SLSB, School of Life Sciences and Biotechnology, Shanghai Jiao Tong University for technical support with fermentation and biodegradation.

## Author Contribution

L.Z. and B.W. conceived the concept of VenusPETase and developed the high-level plans for the project. B.W. and M.L. developed the methodology for VenusPETase design (PET-Flow) with the help from Y.T. and P.T. B.W. and J.L. conducted the enzymatic assays, thermostability assays, and protein crystallization with the help from R.H. and Y.L. L.Z. and B.Zhong analyzed the protein structure. J.Z. performed molecular dynamics simulations with the supervision of W.Q. J.L. performed PET synthesis and injection modeling with the help from X.W. L.Z., J.Z., M.L., and B.W. presented the data. L.Z., B.W., M.L., and J.Z. wrote the manuscript. L.Z. and L.H. supervised the project.

## Additional Information

Correspondence and requests for materials should be addressed to Lirong Zheng or Liang Hong.

## Data Availability

The refined model of VenusPETase in this study has been deposited in the Protein Data Bank with PDB codes 9VJ7. The data of PET-Gym can be downloaded at https://github.com/pet-flow815/petflow.

## Code Availability

The source code of PET-Flow can be found at https://github.com/pet-flow815/petflow.

---

**Extended Data Fig. 1.**
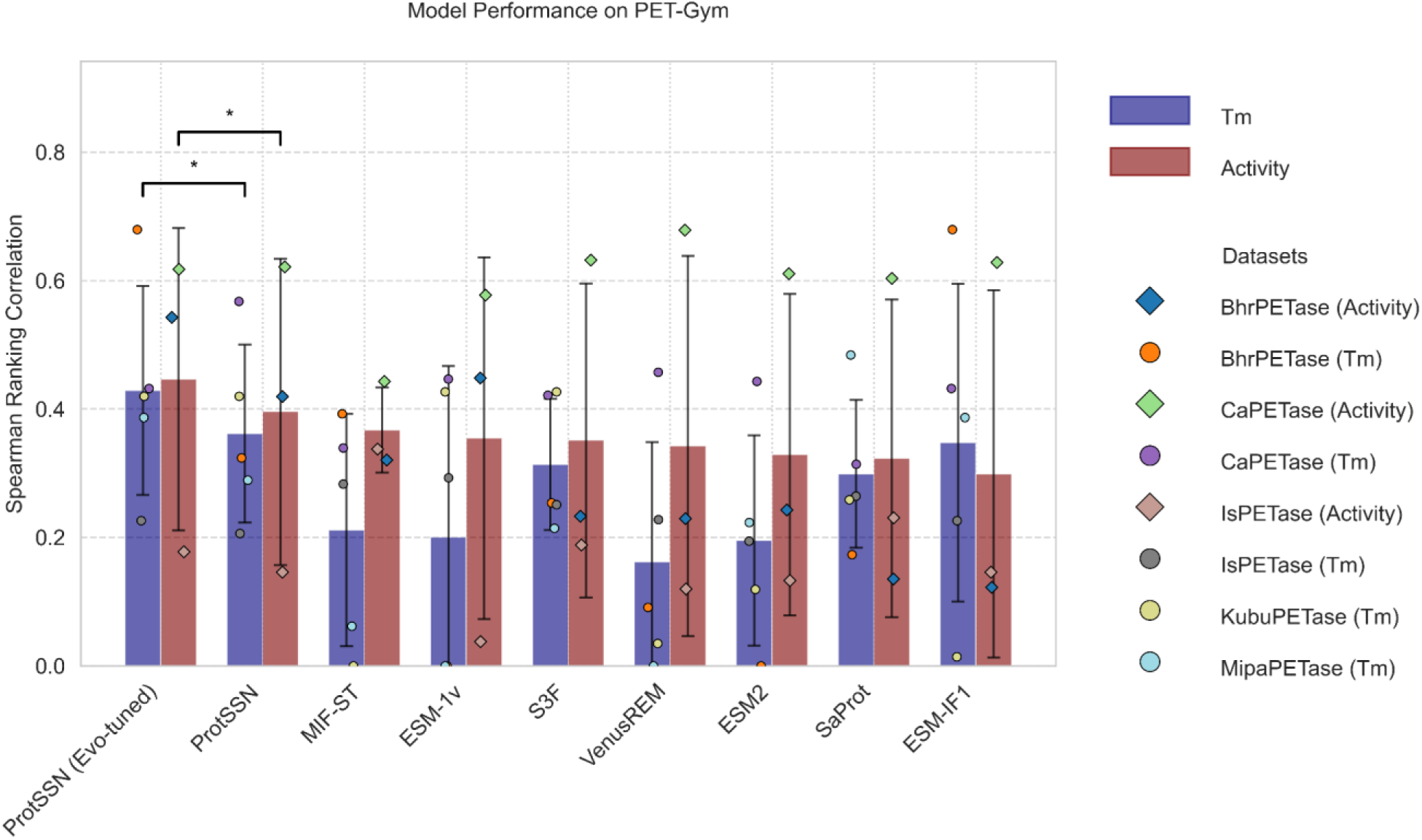
Benchmark results for zero-shot fitness prediction methods. Performance of zero-shot predictors on PET-Gym single-site mutant datasets. For each method, Spearman rank correlation was calculated between predicted mutant fitness scores and experimentally measured melting temperature (Tm, blue bars) or catalytic activity (red bars) across PETase variants. Bars represent the average Spearman correlation across available datasets, and error bars indicate variation across datasets. Colored points denote results for individual enzyme-property datasets, including BhrPETase, CaPETase, IsPETase, KubuPETase, and MipaPETase. Higher Spearman correlation indicates better agreement with experimental mutant effects. ProtSSN with evolutionary fine-tuning showed the strongest overall performance among the evaluated zero-shot methods. Statistical comparisons were performed using two-sided t-tests across datasets. *P < 0.05.

**Extended Data Fig. 2.**
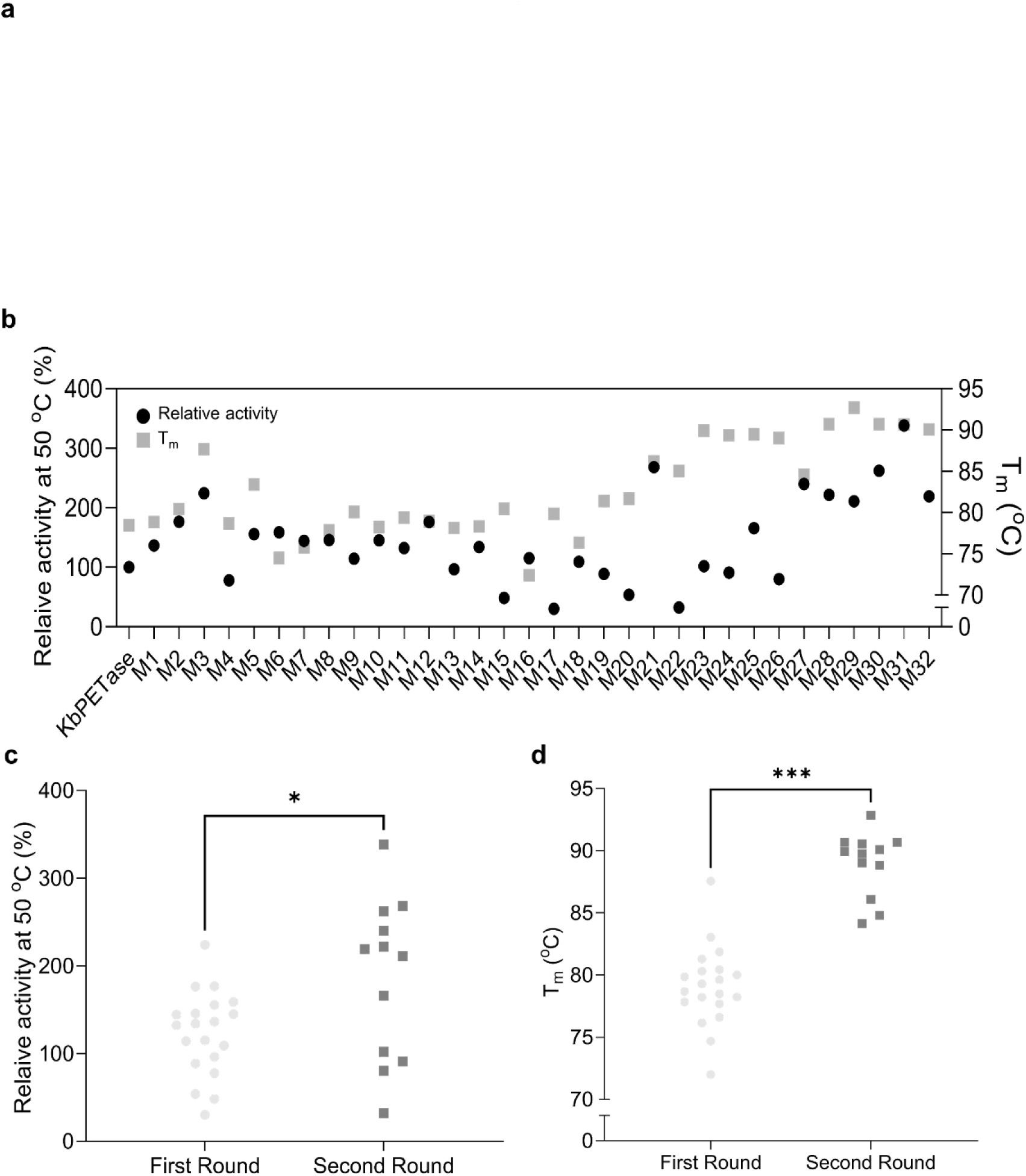
PET-Flow-guided mutagenesis and iterative engineering of VenusPETase leads to enhanced thermostability and catalytic activity. **a**, Structural mapping of single-point mutations selected through PET-Flow-assisted mutational scanning, shown on the VenusPETase model. **b**, Relative PET depolymerization activity at 50 ℃ (black circles, left axis) and melting temperature (T_m_) (gray squares, right axis) of VenusPETase variants carrying single or combinatorial mutations. The activities were homogenized according to KbPETase. **c,** Relative PET-hydrolytic activity at 50 ℃ of enzyme variants from each round of engineering. Activity increases significantly from the first to second round, culminating in variants with >350% activity relative to KbPETase. The activities were homogenized according to KbPETase. **d**, Melting temperature of variants from each engineering round. Second-round VenusPETase variants show substantial increases in thermostability, reaching up to 90.7 °C. PETase thermostability was determined by differential scanning fluorimetry. Statistical comparisons were performed using two-sided unpaired t-tests with Bonferroni correction. *P < 0.05; ***P < 0.001.

**Extended Data Fig. 3.**
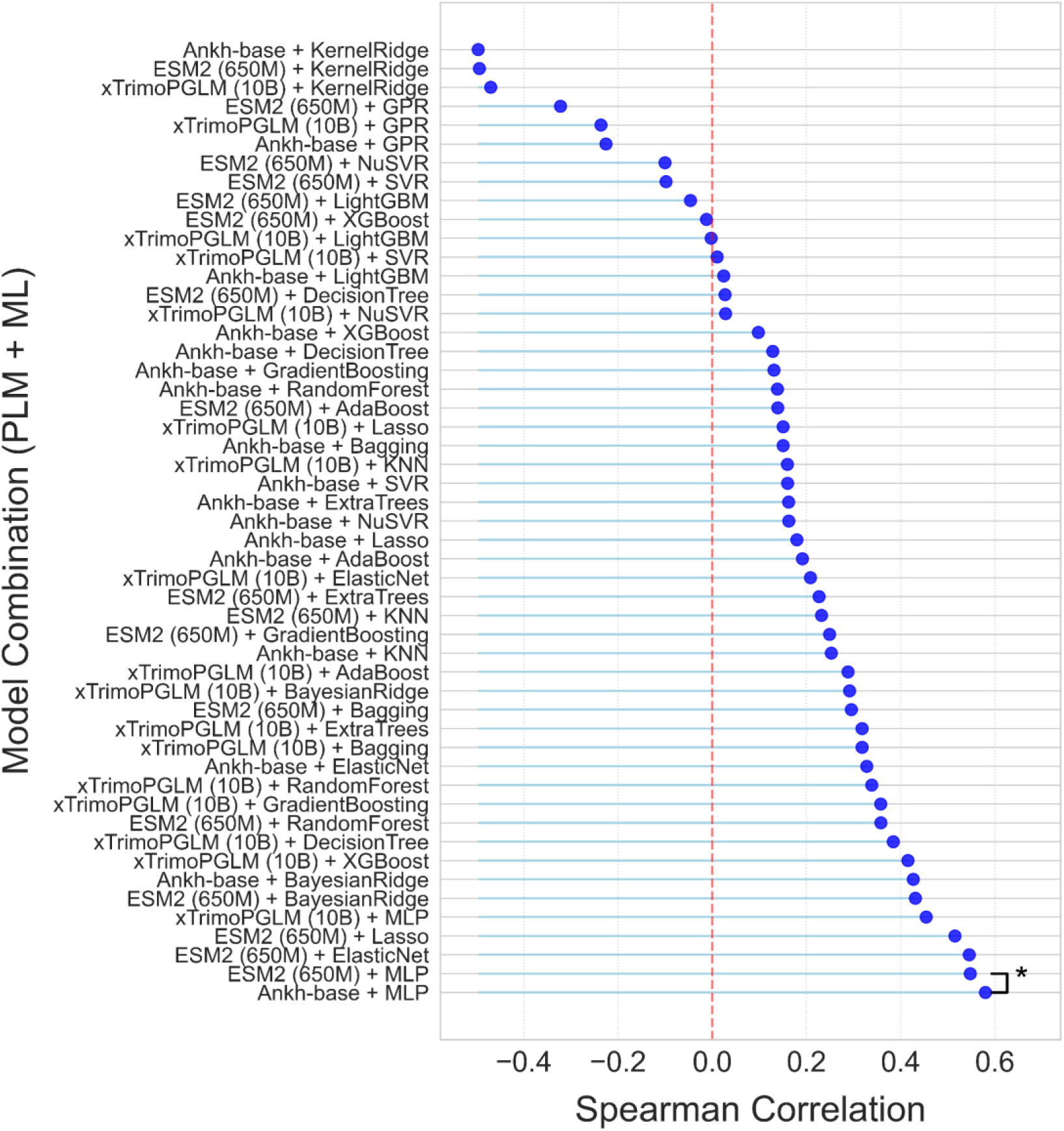
Generalization from single-site to multi-site mutant prediction. Performance of supervised models trained on single-site PETase mutants and evaluated on held-out multi-site mutants in PET-Gym. Each point represents the average Spearman rank correlation between predicted and experimentally measured mutant effects for a protein language model encoder and machine learning regressor combination. Models are ordered by performance from lowest to highest. Among all evaluated combinations, Ankh-base embeddings with an MLP regressor achieved the highest performance, supporting its use for supervised prediction in PET-Flow. Statistical comparisons were performed using two-sided t-tests across datasets. *P < 0.05.

**Extended Data Fig. 4.**
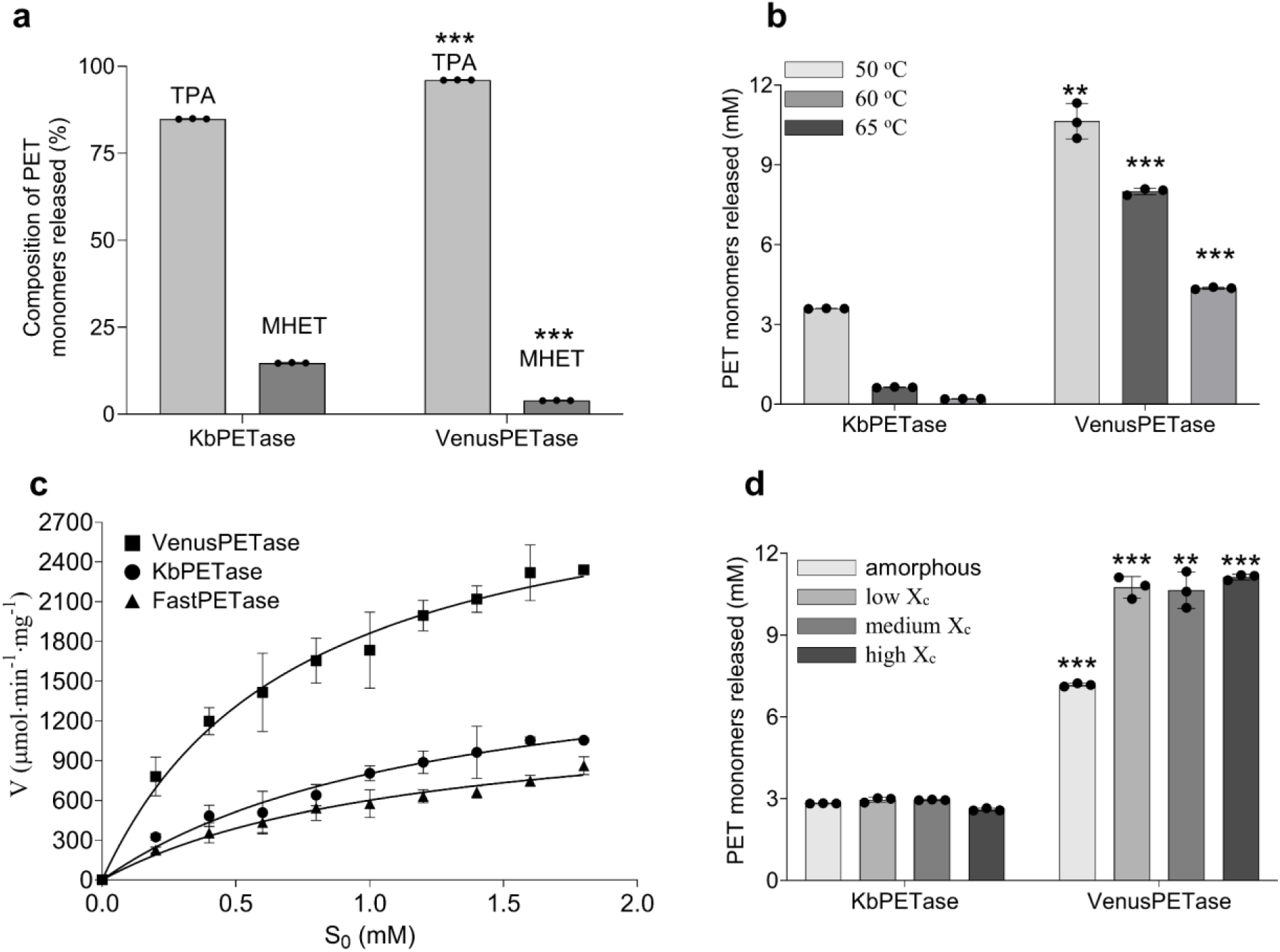
VenusPETase exhibits enhanced product selectivity, catalytic activity, and substrate accessibility across temperatures and PET crystallinity levels. **a**, Composition of PET hydrolysis products released by KbPETase and VenusPETase following 24 h incubation with semi-crystalline PET powder (39.8% crystallinity) at 50 ℃. **b**, Total PET monomer release by KbPETase and VenusPETase at 50, 60, and 65 ℃. Reactions were performed with 0.5 μM enzyme in 100 mM phosphate buffer (pH 8.0) using semi-crystalline PET powder as substrate. **c**, Michaelis–Menten kinetic analysis of VenusPETase, KbPETase, and FastPETase using *p*PNB as a model substrate. **d**, PET monomer release from amorphous PET (Xc ≈ 7.9%), low-crystallinity (Xc ≈ 12.0%), medium-crystallinity (Xc ≈ 32.9 %), and high-crystallinity (Xc ≈ 46.3%) PET substrates after 24 h enzymatic treatment at 50 ℃ with VenusPETase or KbPETase. All data represent mean ± s.d. From three independent biological replicates (n = 3). Statistical significance was determined by one-sided unpaired t-tests. **P < 0.01; ***P < 0.001.

**Extended Data Fig. 5.**
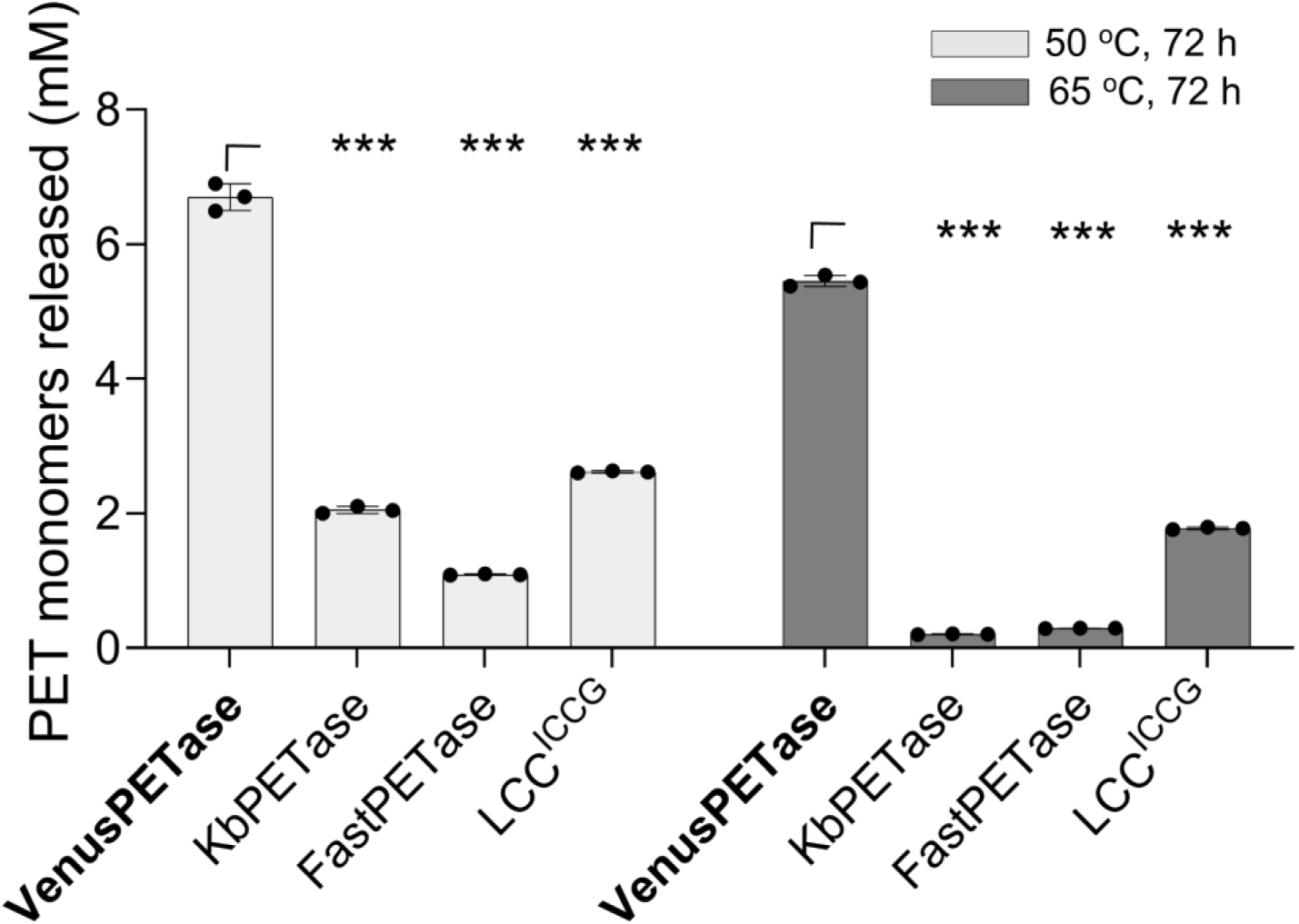
Hydrolytic activities of VenusPETase, KbPETase, FastPETase, and LCC ^ICCG^ after thermal treatment. The activities of VenusPETase, KbPETase, FastPETase, and LCC^ICCG^ were measured after thermal treatment at 50 °C or 65 °C for 3 days. Following thermal treatment, each PETase was assayed under its individual optimal catalytic temperature and buffer conditions. All data represent mean ± s.d. from three independent biological replicates (n = 3). Statistical analysis was performed using two-way ANOVA followed by Dunnett’s multiple comparisons test. **P < 0.01, ***P < 0.001.

**Extended Data Fig. 6.**
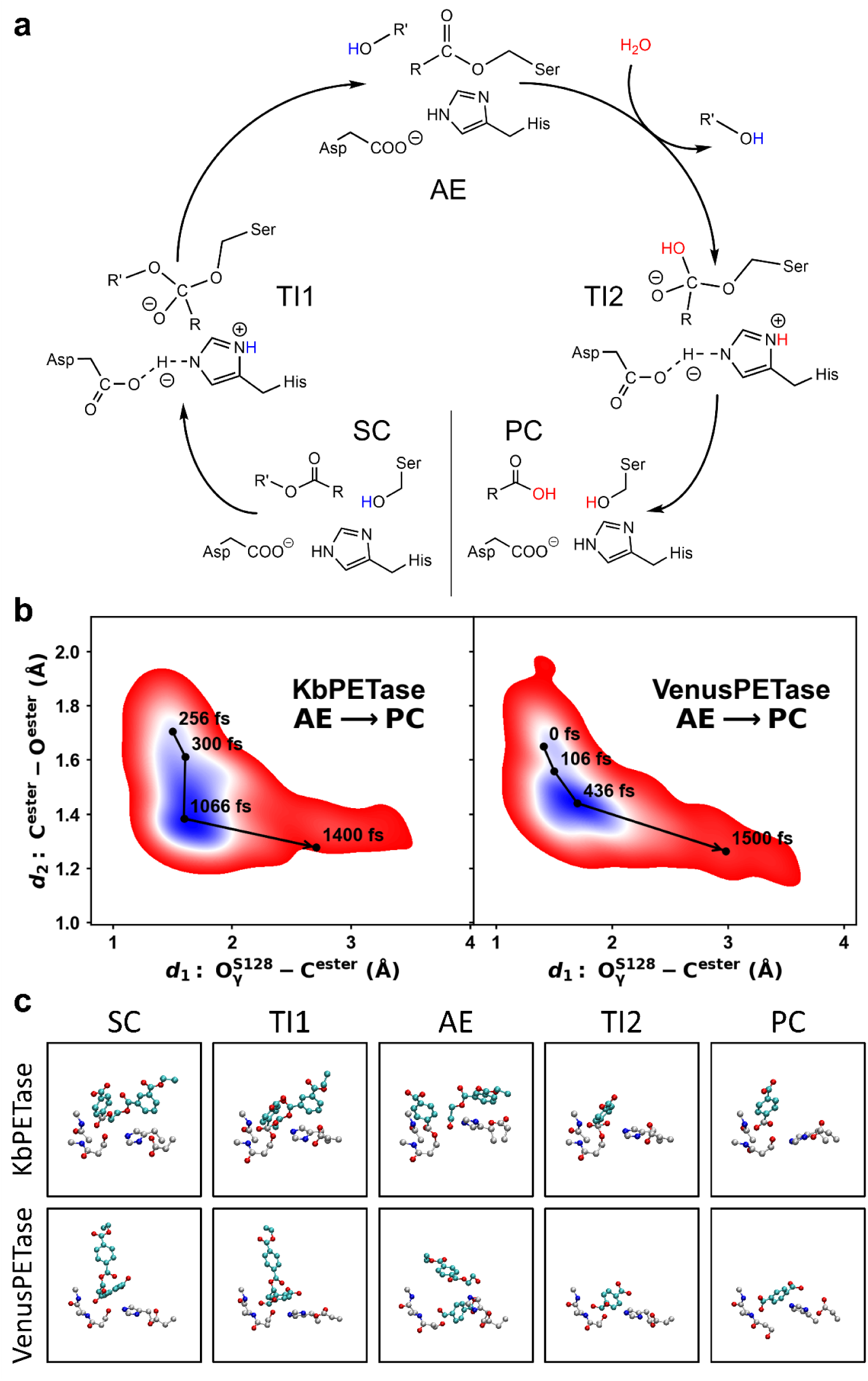
Structural analysis of the constructed CPET and APET structures. **a**, Mechanism of PETase catalyzing ester bond cleavage in PET *via* an acylation-deacylation mechanism. In the acylation phase (SC→TI1→AE), the PET chain binds to the enzyme’s active site, forming a substrate complex (SC). The catalytic serine is activated by proton transfer to histidine, stabilized by aspartic acid, and attacks the carbonyl carbon, forming a tetrahedral intermediate (TI1) stabilized by an oxyanion hole. Then cleavage of the C–O bond releases ethylene glycol and forms the acyl-enzyme intermediate (AE). Next, in the deacylation phase (AE→TI2→PC), a water molecule, activated by histidine, attacks the AE carbonyl, forming a second tetrahedral intermediate (TI2). Then the acyl-enzyme bond breaks, releasing terephthalic acid and regenerating free enzymes. Finally, the histidine is deprotonated and returns to the initial state, ready for the next catalytic cycle. **b**, Key distances change among metadynamics trajectory, which are related to the catalysis process deacylation phase. Among meta-FPMD simulations, VenusPETase showed smaller energy barriers than KbPETase during the reaction process, indicating higher catalytic ability of PET degradation. **c**, Snapshots of the key conformations of the states in **a** during degradation processes. The carbons of substrate are colored with cyan, and carbons of enzyme residues are colored with gray.

**Extended Data Fig. 7.**
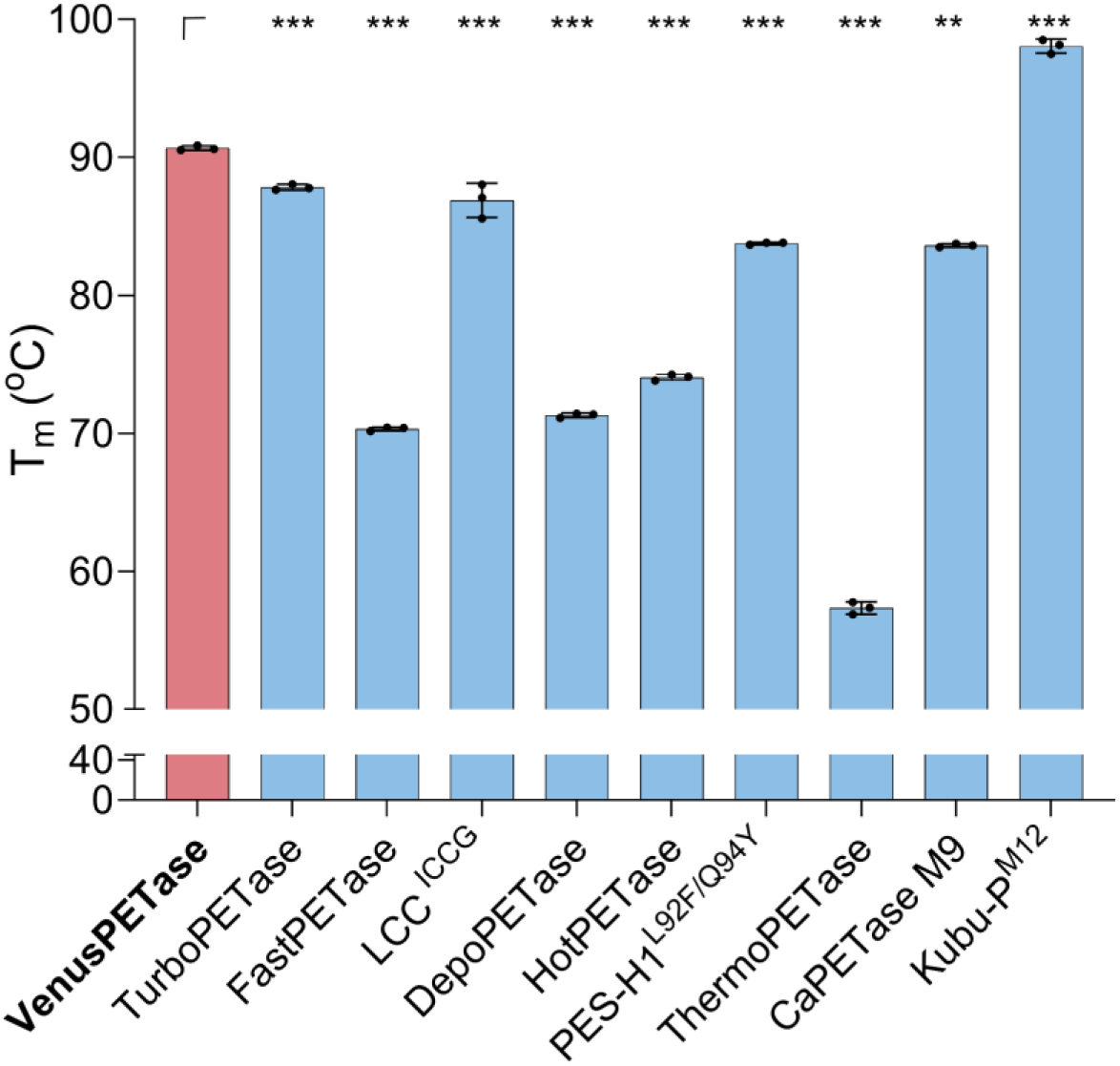
Melting temperature of VenusPETase and other PETase variants determined by differential scanning fluorimetry. All data represent mean ± s.d. from three independent biological replicates (n = 3). Statistical comparisons were performed using one-sided unpaired t-tests. **P < 0.01; ***P < 0.001.

**Extended Data Fig. 8.**
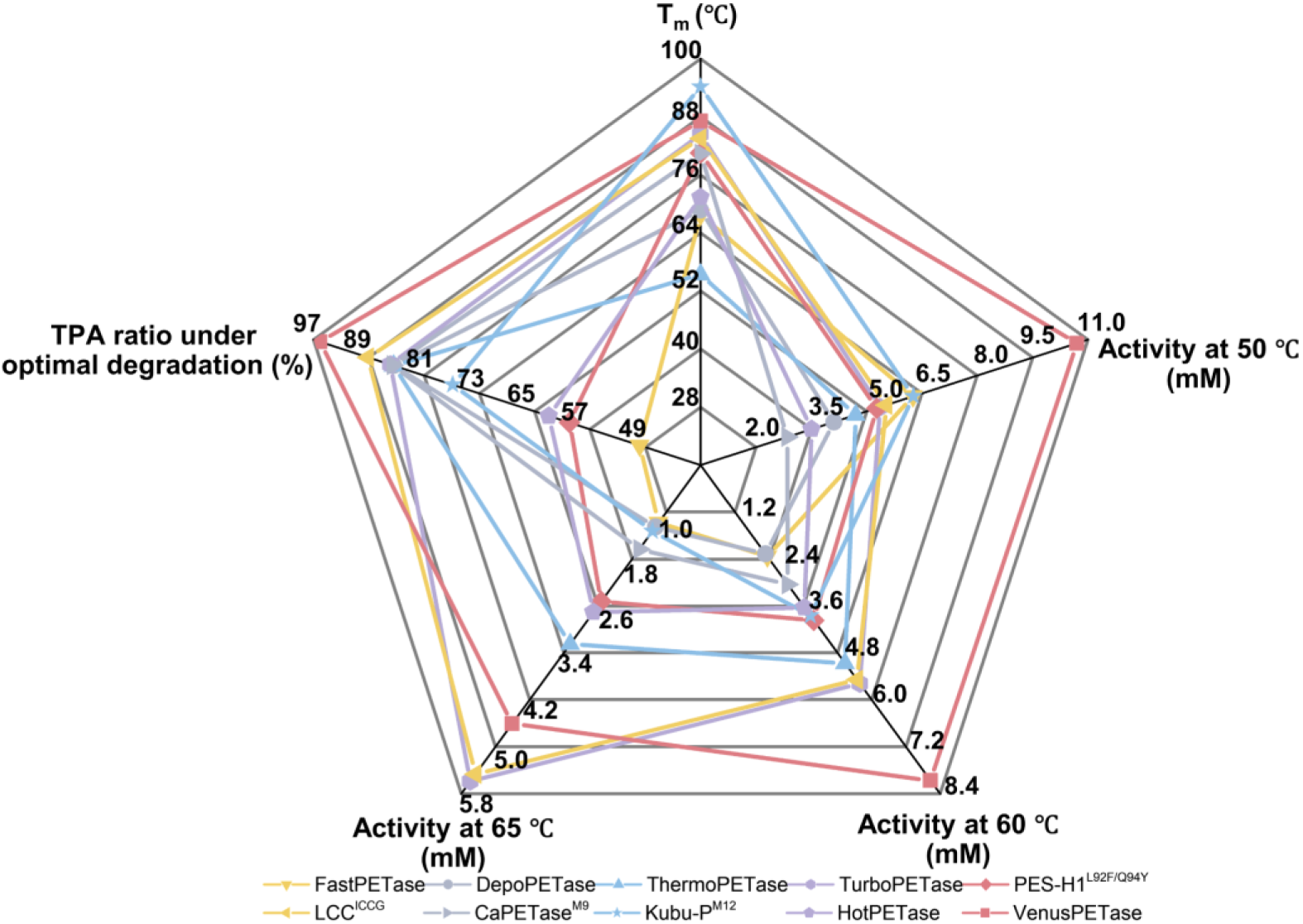
Multidimensional performance comparison of VenusPETase and other PETase variants across thermostability and hydrolytic metrics. Radar plot showing the performance of VenusPETase and nine PETase variants across five properties. From the top in clockwise order, the axes correspond to T_m_, hydrolytic activity at 50 °C, hydrolytic activity at 60 °C, hydrolytic activity at 65 °C, and the release of TPA under the respective optimal reaction conditions. Each colored polygon represents a distinct PETase variant or VenusPETase.

**Extended Data Fig. 9.**
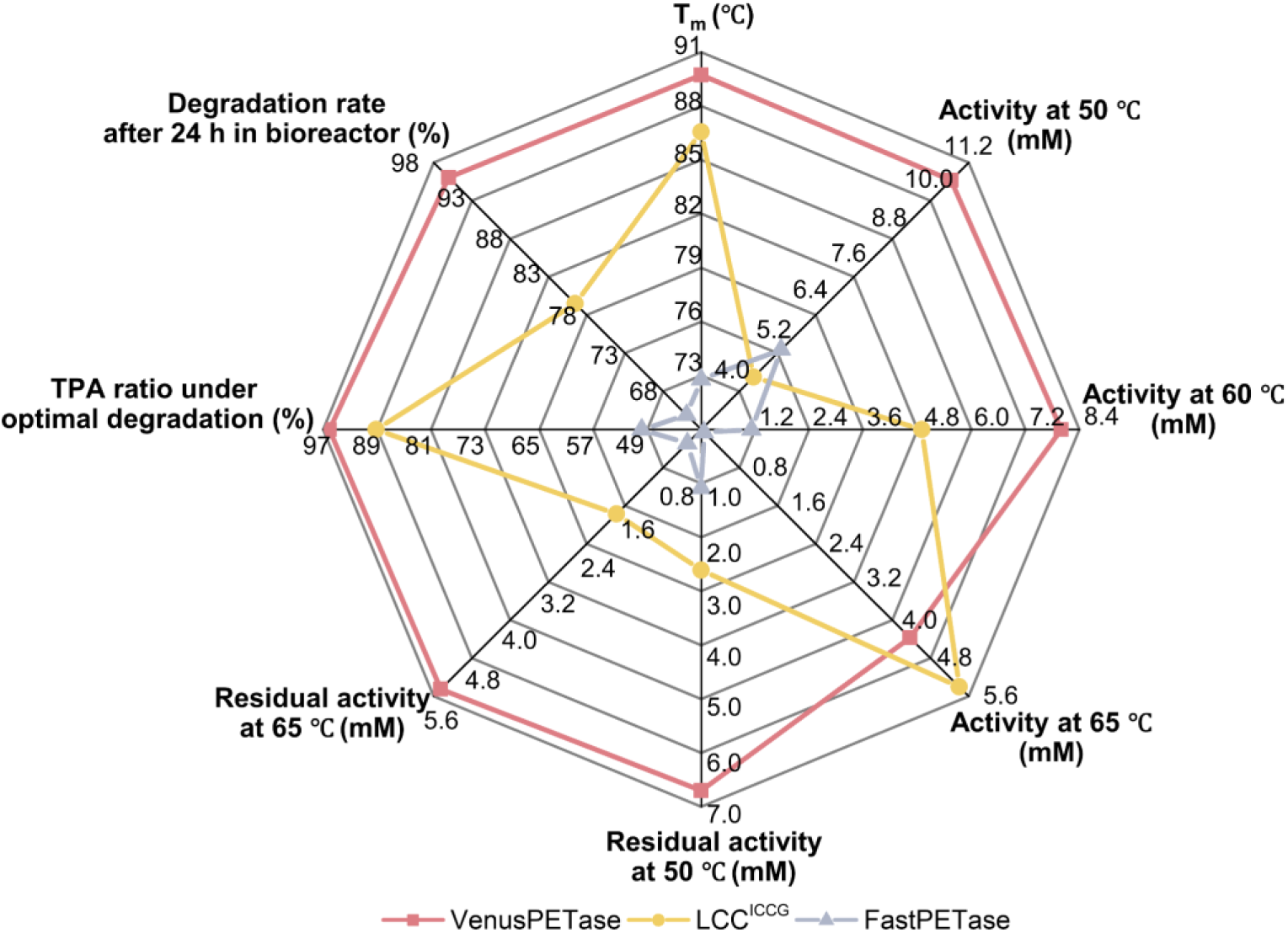
Multidimensional performance comparison of VenusPETase, LCC^ICCG^, and FastPETase. Radar plot comparing the overall performance of VenusPETase, LCC^ICCG^, and FastPETase across eight properties. The axes represent melting temperature, hydrolytic activity at 50 °C, hydrolytic activity at 60 °C, hydrolytic activity at 65 °C, hydrolytic activity after heat treatment at 50 °C, hydrolytic activity after heat treatment at 65 °C, the release of TPA under the respective optimal reaction conditions, and the 24 h degradation rate in a 5 L bioreactor. Red, yellow, and gray traces denote VenusPETase, LCC^ICCG^, and FastPETase, respectively.

**Extended Data Table 1.** Kinetic parameters of KbPETase, VenusPETase, and FastPETase derived from the Michaelis-Menten experiments at 50 °C.

| PETase | $K_M$ (mM) | $k_{cat}$ ($s^{-1}$) | $k_{cat}/K_M$ ($mM \cdot s^{-1}$) |
| --- | --- | --- | --- |
| KbPETase | $1.04 \pm 0.201$ | $0.270 \pm 0.021$ | $0.263 \pm 0.031$ |
| FastPETase | $1.57 \pm 0.416$ | $0.215 \pm 0.029$ | $0.141 \pm 0.02$ |
| VenusPETase | $0.539 \pm 0.023$ | $0.508 \pm 0.0172$ | $0.539 \pm 0.023$ |

**Extended Data Table 2.**
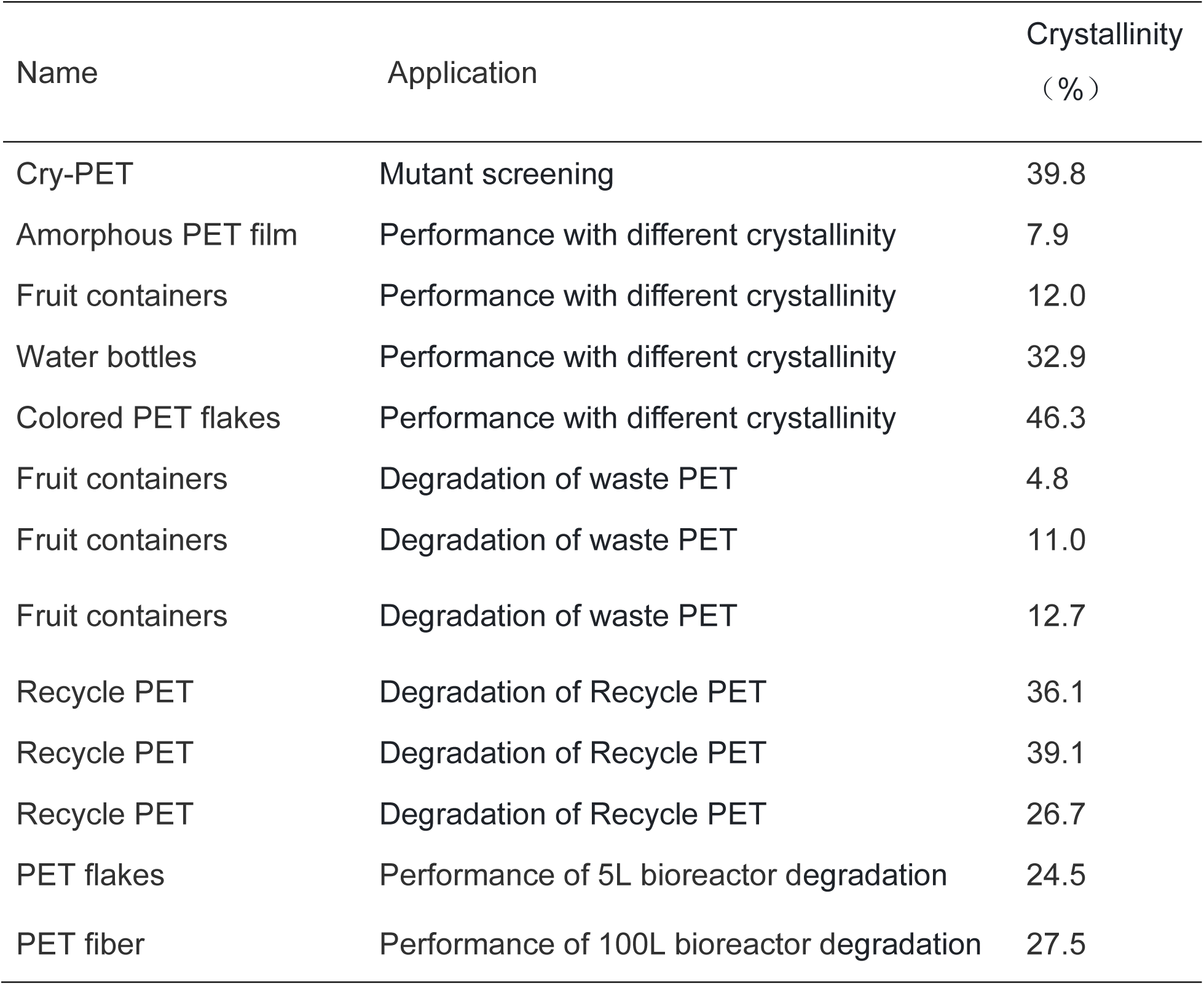
PET substrates used in this study and crystallinity.

| Name | Application | Crystallinity<br>(%) |
| --- | --- | --- |
| Cry-PET | Mutant screening | 39.8 |
| Amorphous PET film | Performance with different crystallinity | 7.9 |
| Fruit containers | Performance with different crystallinity | 12.0 |
| Water bottles | Performance with different crystallinity | 32.9 |
| Colored PET flakes | Performance with different crystallinity | 46.3 |
| Fruit containers | Degradation of waste PET | 4.8 |
| Fruit containers | Degradation of waste PET | 11.0 |
| Fruit containers | Degradation of waste PET | 12.7 |
| Recycle PET | Degradation of Recycle PET | 36.1 |
| Recycle PET | Degradation of Recycle PET | 39.1 |
| Recycle PET | Degradation of Recycle PET | 26.7 |
| PET flakes | Performance of 5L bioreactor degradation | 24.5 |
| PET fiber | Performance of 100L bioreactor degradation | 27.5 |

## Supplementary Information

**Supplementary Figure 1.**
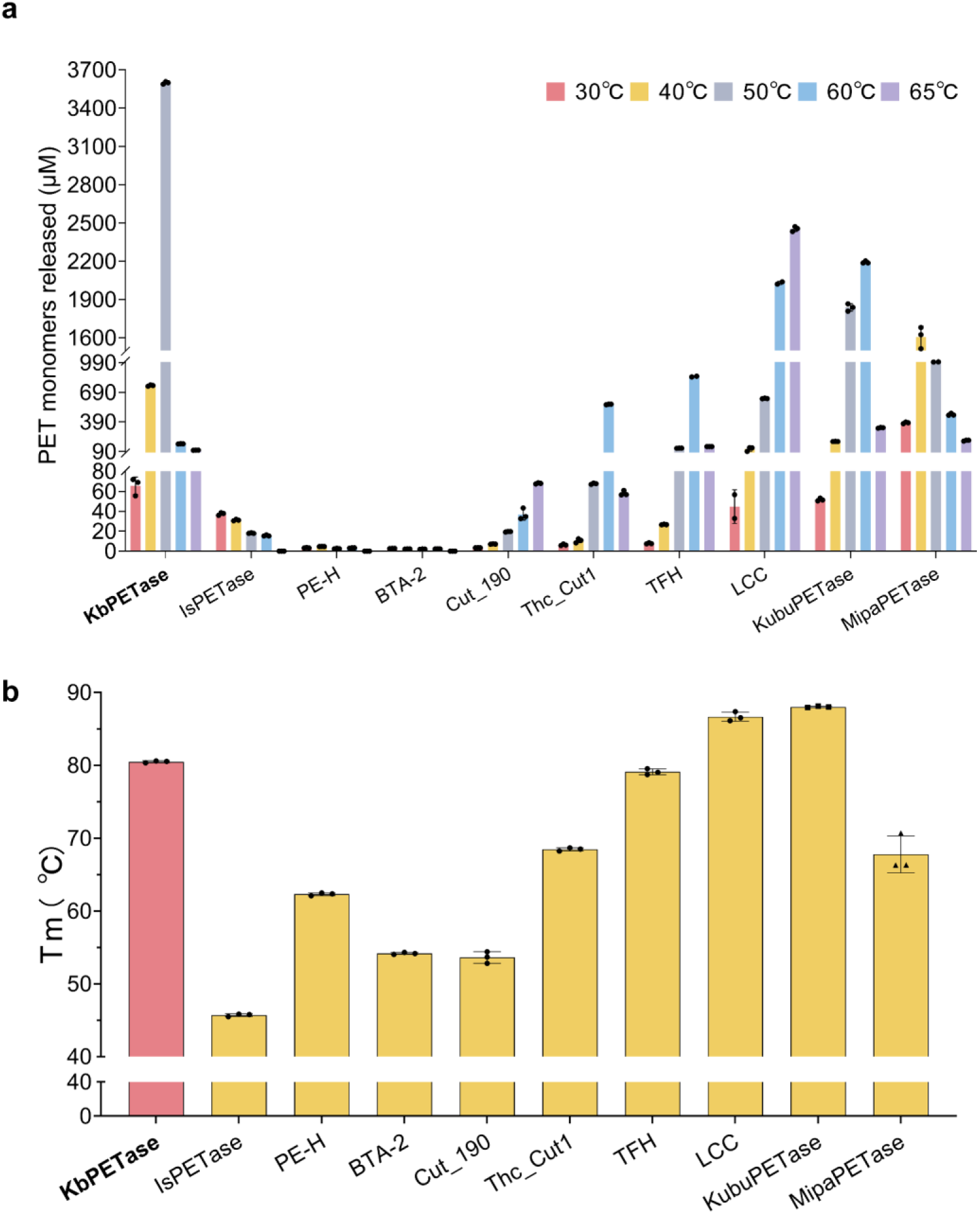
Performance comparison of wild-type PETases. **a**, Release of PET monomers from PET film hydrolysis by different wild-type PET hydrolases at 30–65 °C for 72 h, with a solids loading of 15 g kg⁻¹ and an enzyme loading of 3 mg_enzyme_ g_PET_⁻¹, in 50 mM glycine–NaOH buffer (pH 9.0). **b**, Thermostability comparison of different wild-type PETases. T_m_ values were determined by DSF.

**Supplementary Figure 2. Differential scanning fluorimetry curves of mutants obtained during evolution.** The intersection of the black arrow line with the X-axis represents the T_m_ value.

**Supplementary Figure 3.**
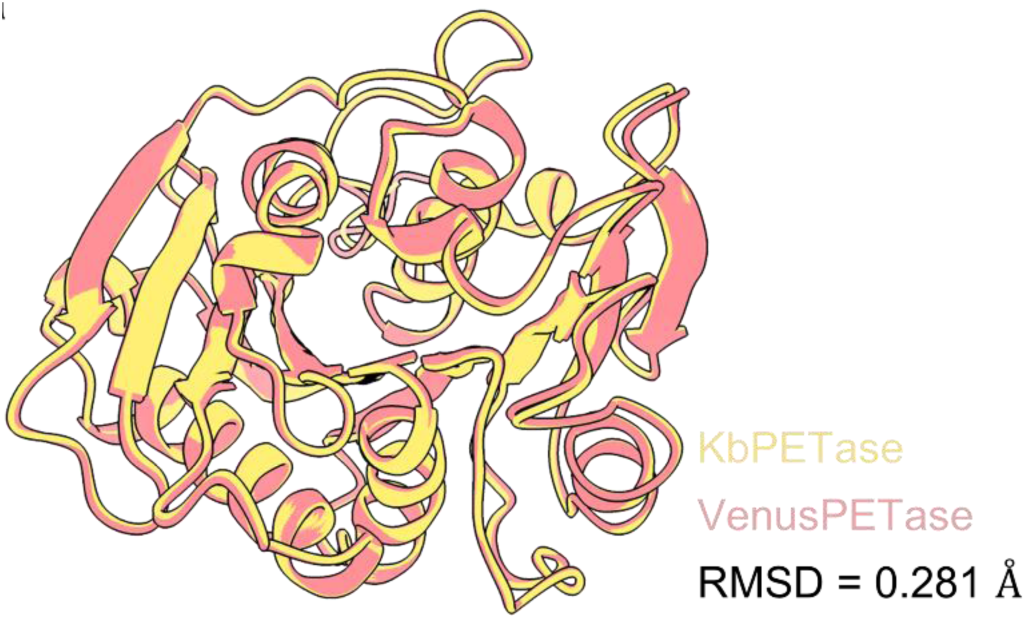
Structural comparison between VenusPETase and KbPETase. Superposition of the overall structures of VenusPETase and KbPETase using PyMOL.

**Supplementary Figure 4.**
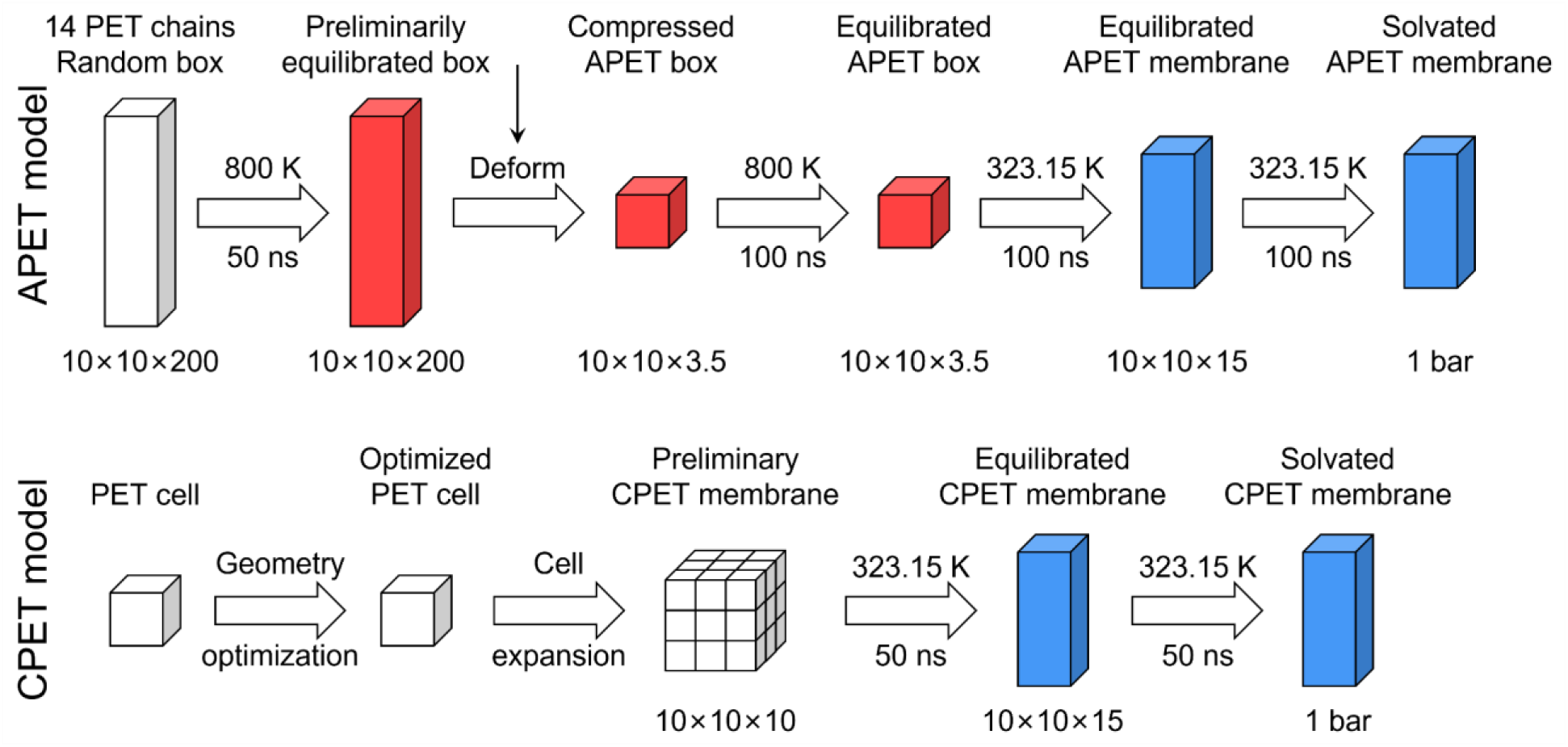
Structure construction protocol of low-crystallinity PET (APET) and high-crystallinity PET (CPET). The APET structure was constructed by compressing 14 PET chains (each contains 100 PET residues) into a 10 × 10 × 3.5 nm box, and then solvated in a 10 × 10 × 15 nm box to form membrane structure after equilibration. The CPET cell was optimized according to crystal structure and then expanded to a super cell with similar size as APET membrane, and then solvated in a 10 × 10 × 15 nm box.

**Supplementary Figure 5.**
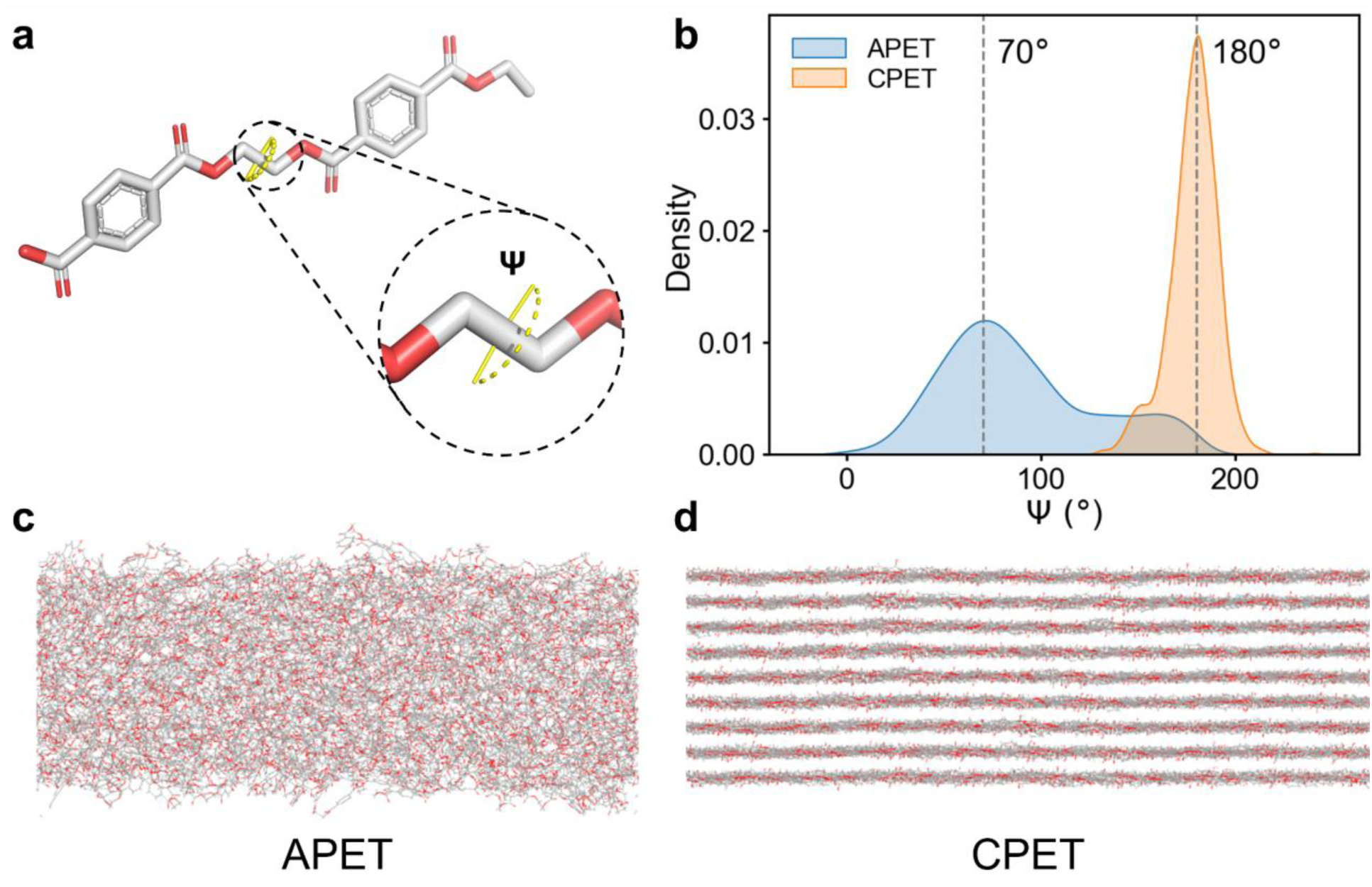
Structural analysis of the constructed CPET and APET structures. **a**, The O–CH_2_–CH_2_–O dihedral (Ψ) could reveal the PET crystallinity, where 180° corresponds to high crystallinity, and 70° corresponds to low crystallinity. **b**, The distribution of Ψ among the two simulation PET structures after 50 ns MD simulation shows consistent crystallinity. **c**,**d**, The equilibrated structures of the simulated **c** APET and **d** CPET structures.

**Supplementary Figure 6.**
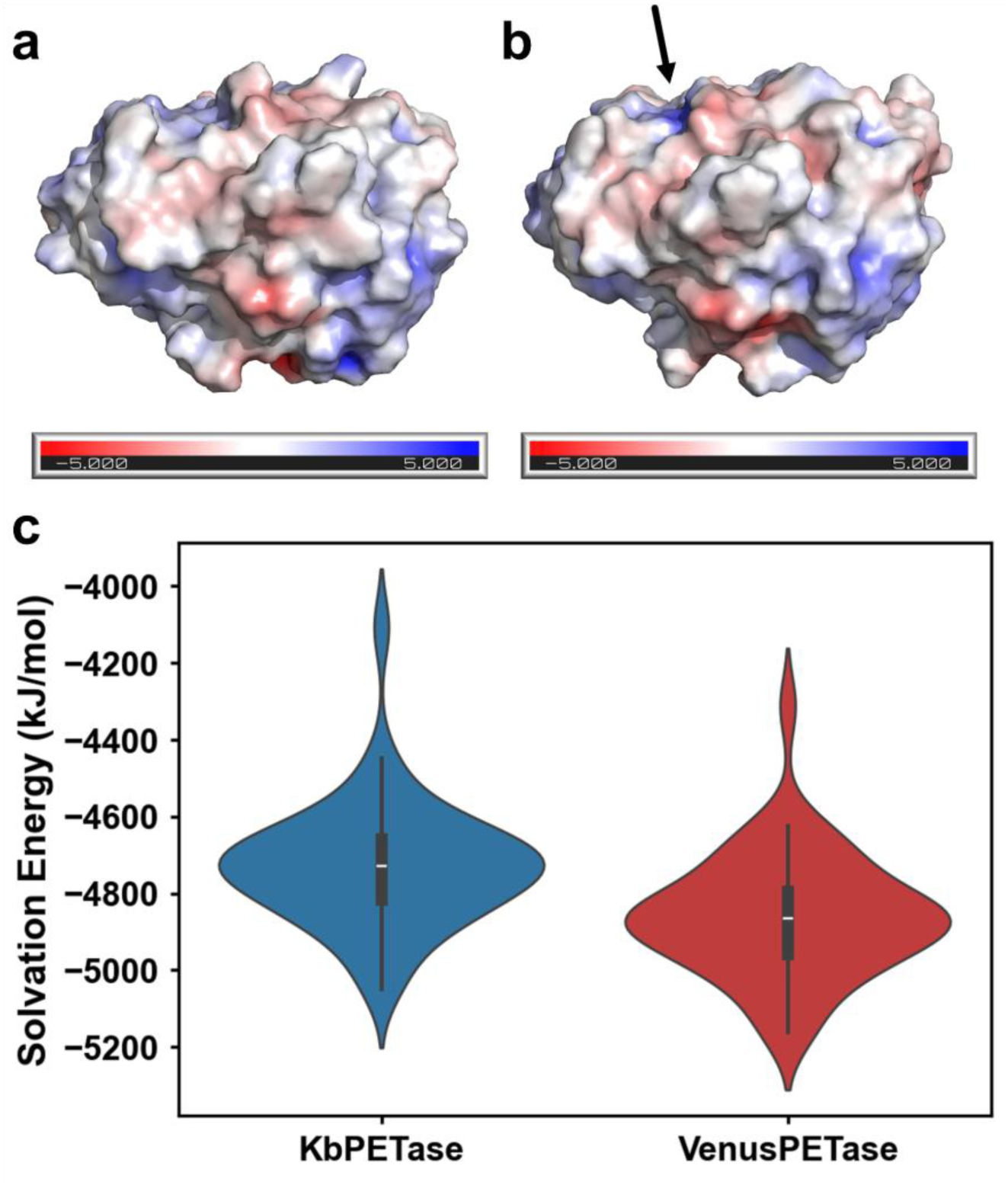
Surface electrostatic potential and solvation energy analysis of the enzymes. **a**,**b**, The surface electrostatic potential of **a** KbPETase and **b** VenusPETase. The VenusPETase showed enhanced potential (arrow) and generated stronger interaction with the PET surface. **c**, The solvation energy of the enzymes. VenusPETase showed a significantly enhanced solvation effect compared to KbPETase.

**Supplementary Figure 7.**
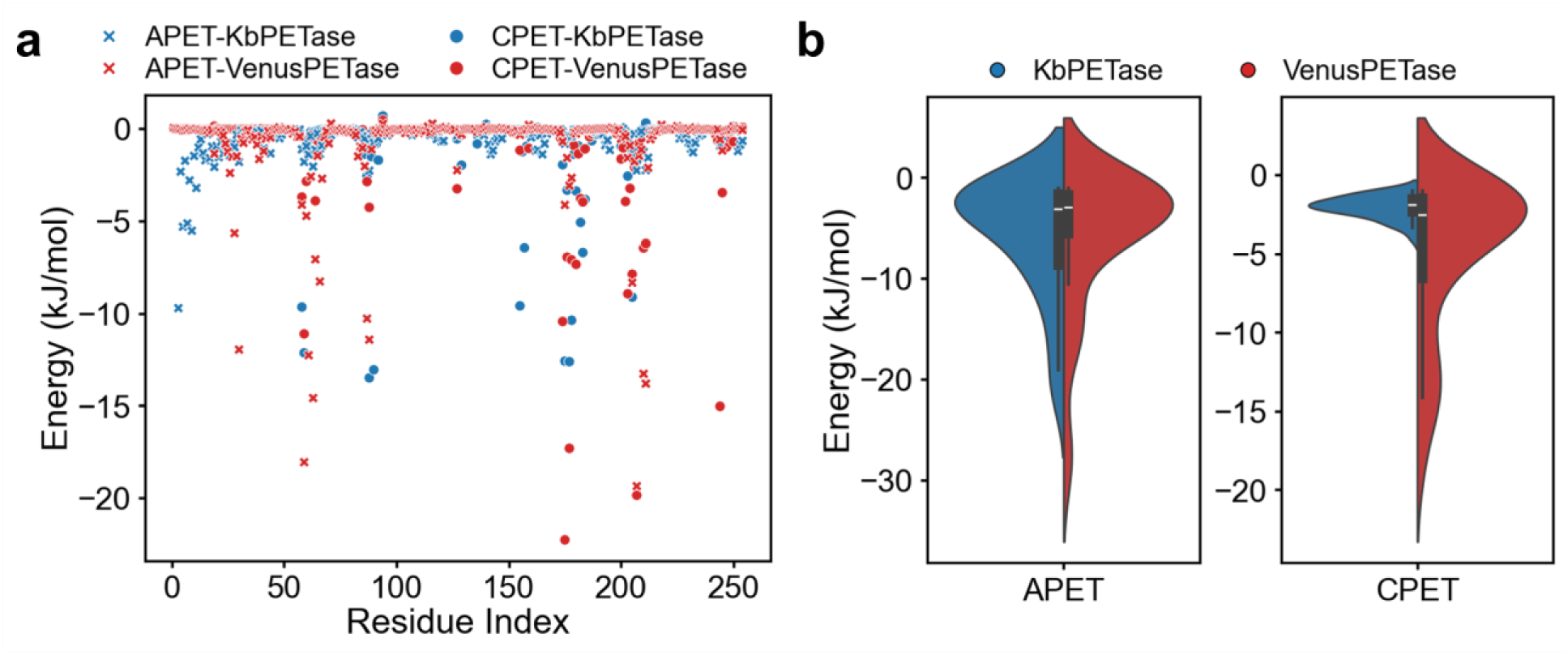
Binding energy analysis of KbPETase and VenusPETase on APET and CPET. **a**, Binding energy residue-decomposition of surface electrostatic potential of **a** KbPETase and **b** VenusPETase. The VenusPETase showed enhanced potential (arrow) and generated stronger interaction with the PET surface. **c**, The solvation energy of the enzymes. VenusPETase showed a significantly enhanced solvation effect compared to KbPETase.

**Supplementary Figure 8.**
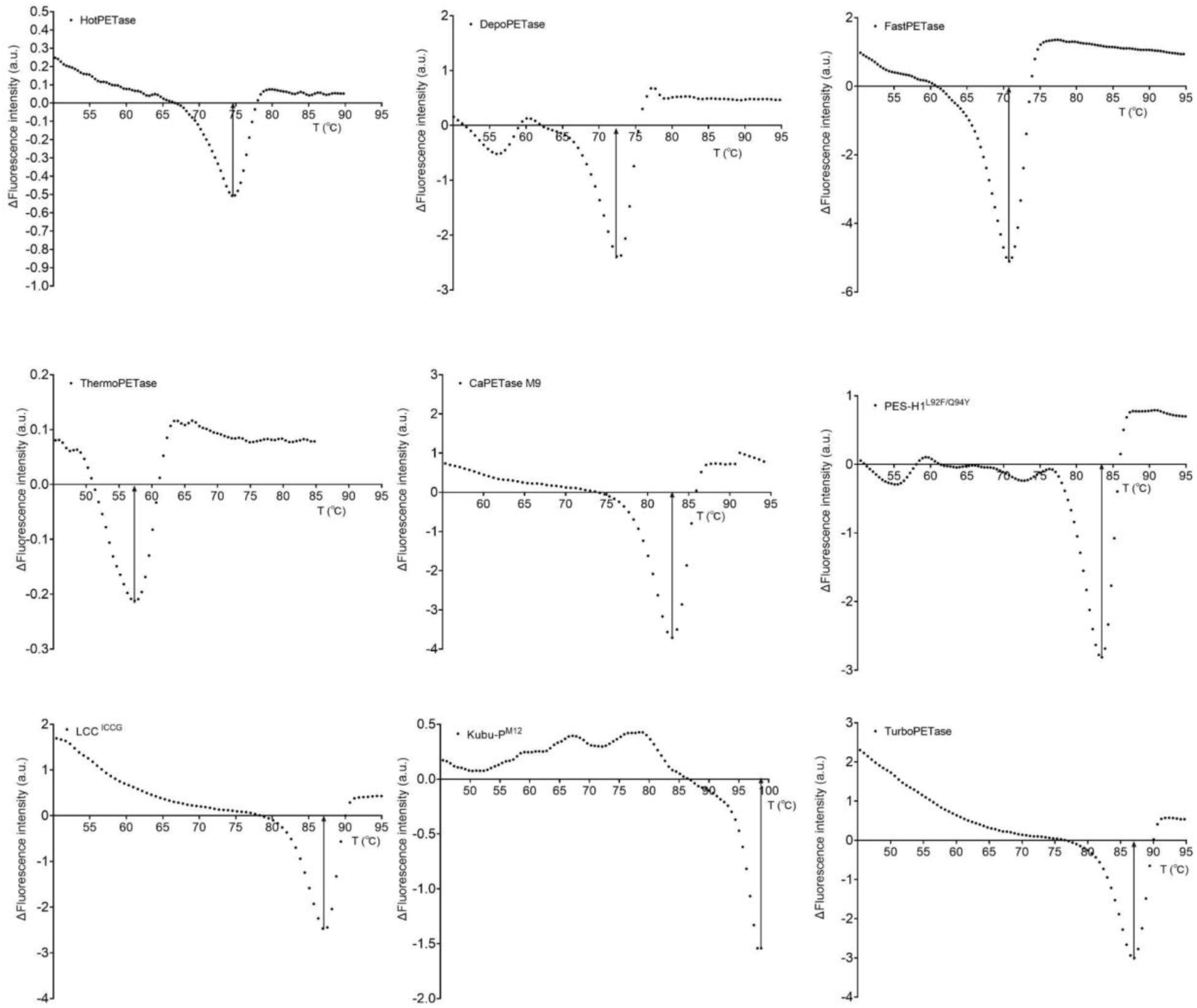
Melting temperature results of other reported PETases measured by differential scanning fluorimetry. The intersection of the black arrow line with the X-axis represents the T_m_ value.

**Supplementary Figure 9.**
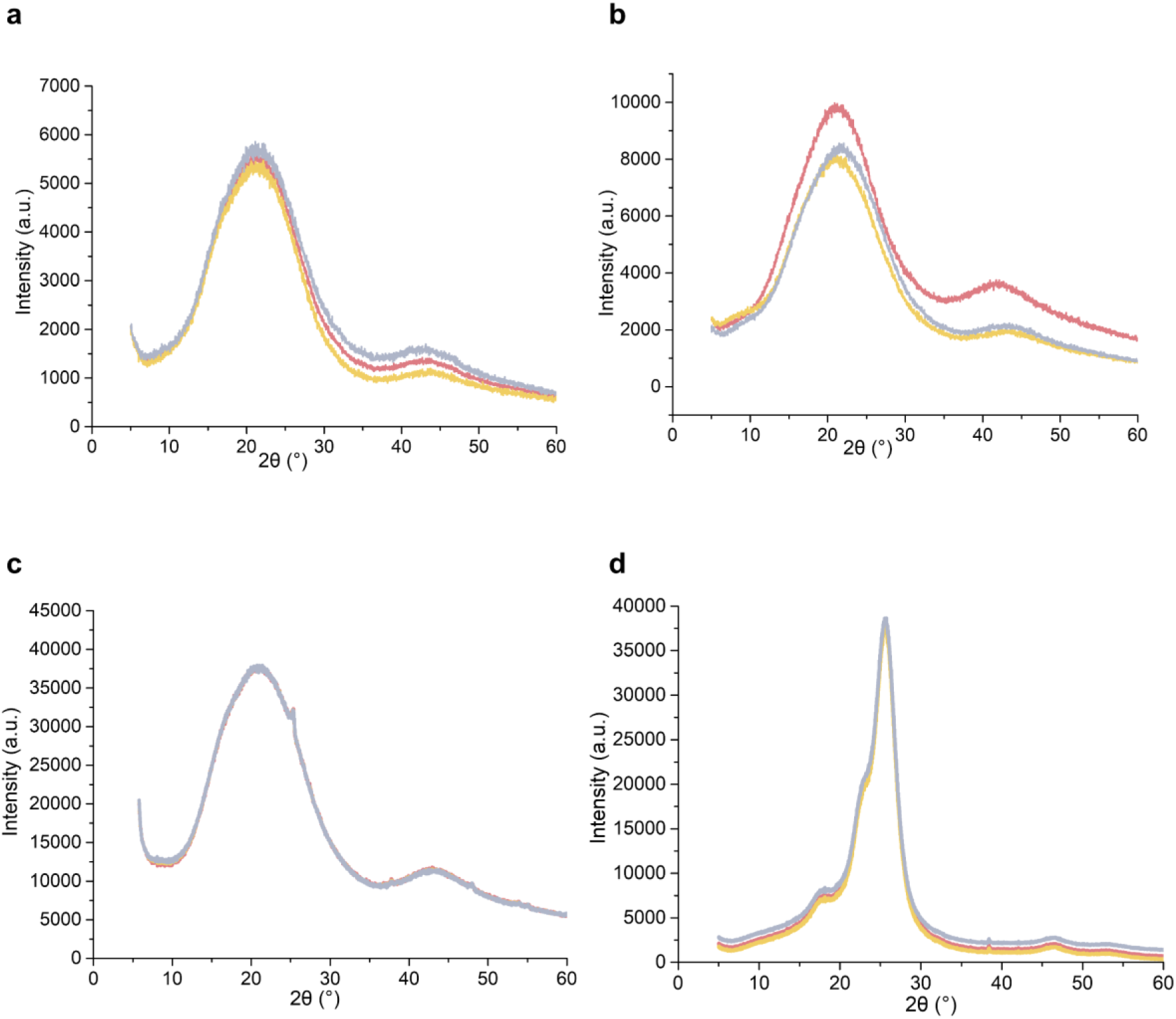
X-ray diffraction of PET substrates with different crystallinities used in this study. **a**, amorphous PET substrate (X_c_ ≈ 7.9%); **b,** low-crystallinity PET substrate (X_c_ ≈ 12.0%); **c**, medium-crystallinity PET substrate (X_c_ ≈ 32.9%); **d**, high-crystallinity PET substrate (X_c_ ≈ 46.3%).

**Supplementary Figure 10.**
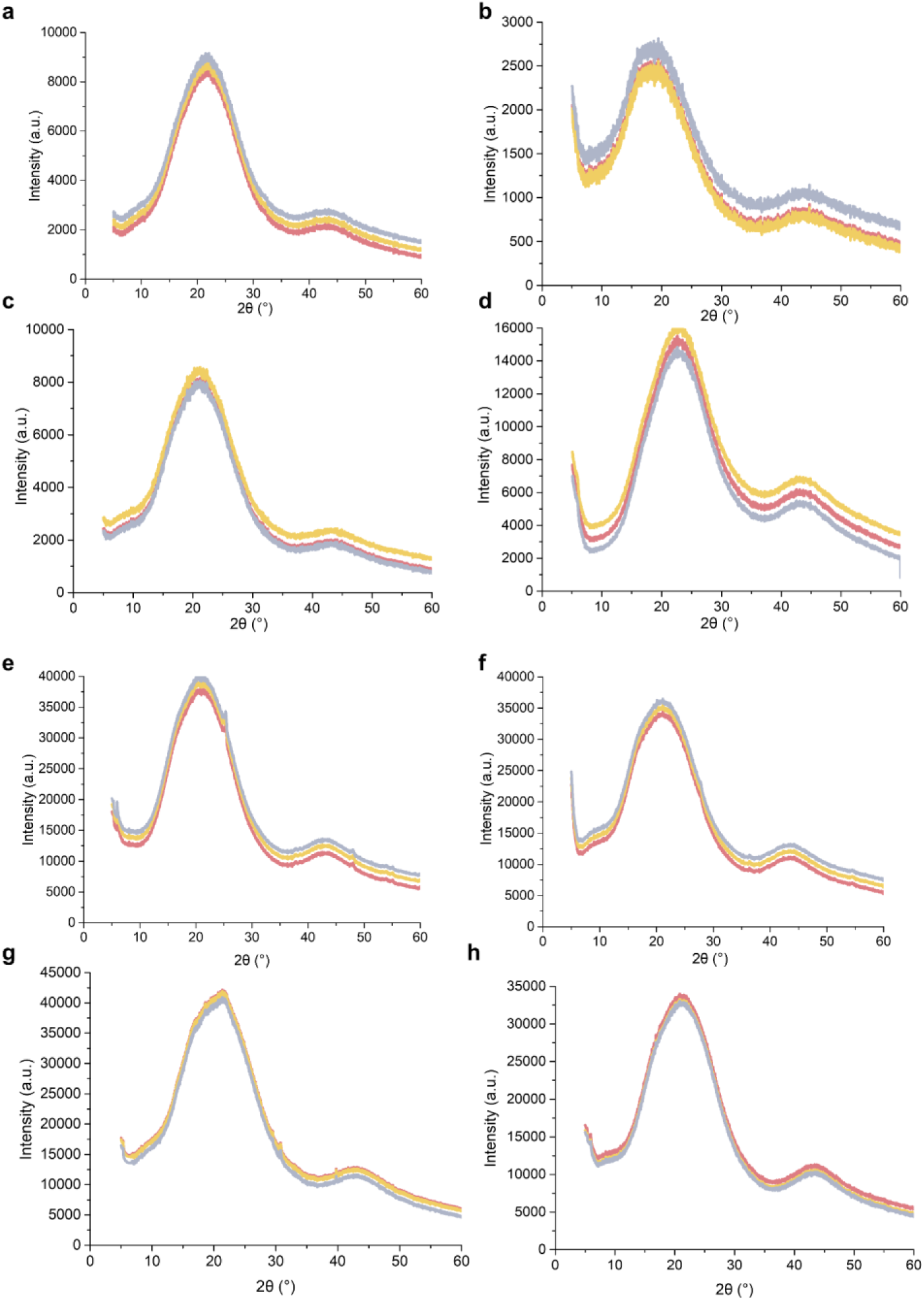
X-ray diffraction (XRD) patterns for crystallinity analysis of post-consumer and recycled PET substrates. **a–c,** samples collected from different locations of a disposable fruit packaging container; **d–f,** recycled PET samples; **g,** PET substrate used in the 5L bioreactor; **h,** PET substrate used in the 100L bioreactor.

**Supplementary Figure 11.**
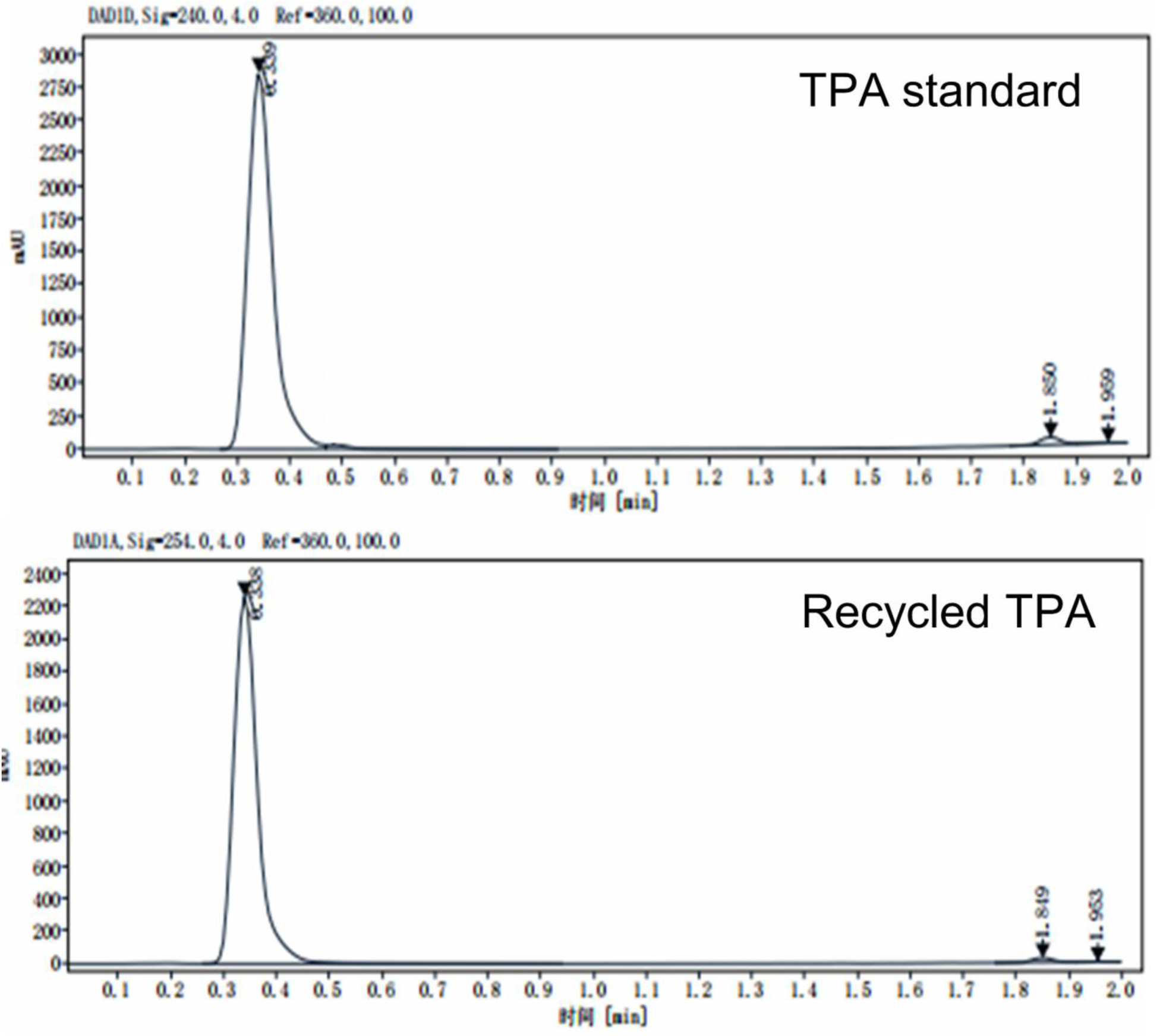
The purity of recycle TPA (rTPA) by UPLC.

**Supplementary Table 1.** Round 1 single-site mutants selected by zero-shot prediction with experimental measurements.

| **Mutant Number** | **T<sub>m</sub> (°C)** | **Activity at 50 °C (%)** |
| --- | --- | --- |
| M1 | 78.86 | 136.40 |
| M2 | 80.41 | 176.73 |
| M3 | 87.65 | 224.17 |
| M4 | 78.68 | 77.88 |
| M5 | 83.38 | 155.52 |
| M6 | 74.48 | 158.91 |
| M7 | 75.77 | 144.41 |
| M8 | 77.84 | 145.64 |
| M9 | 80.08 | 114.31 |
| M10 | 78.22 | 145.06 |
| M11 | 79.38 | 132.20 |
| M12 | 78.94 | 176.33 |
| M13 | 78.13 | 96.44 |
| M14 | 78.30 | 134.07 |
| M15 | 80.46 | 48.29 |
| M16 | 72.39 | 115.17 |
| M17 | 79.84 | 30.26 |
| M18 | 76.35 | 109.18 |
| M19 | 81.38 | 88.81 |
| M20 | 81.67 | 53.82 |

**Supplementary Table 2.** Round 2 multi-site mutants selected by supervised model-guided optimization with experimental measurements. The final optimized variant, VenusPETase, is shown in bold.

| **Mutant Number** | **T<sub>m</sub> (°C)** | **Activity at 50 °C (%)** |
| --- | --- | --- |
| M21 | 86.21 | 268.16 |
| M22 | 85.02 | 32.20 |
| M23 | 89.89 | 101.80 |
| M24 | 89.34 | 91.07 |
| M25 | 89.46 | 165.80 |
| M26 | 89.02 | 80.27 |
| M27 | 84.57 | 240.17 |
| M28 | 90.70 | 221.70 |
| M29 | 92.70 | 211.05 |
| M30 | 90.70 | 262.19 |
| **M31** | **90.68** | **338.15** |
| M32 | 90.07 | 219.08 |

**Supplementary Table 3.** Training and validation set assignments for Round 1 mutants.

**Supplementary Table 4.**
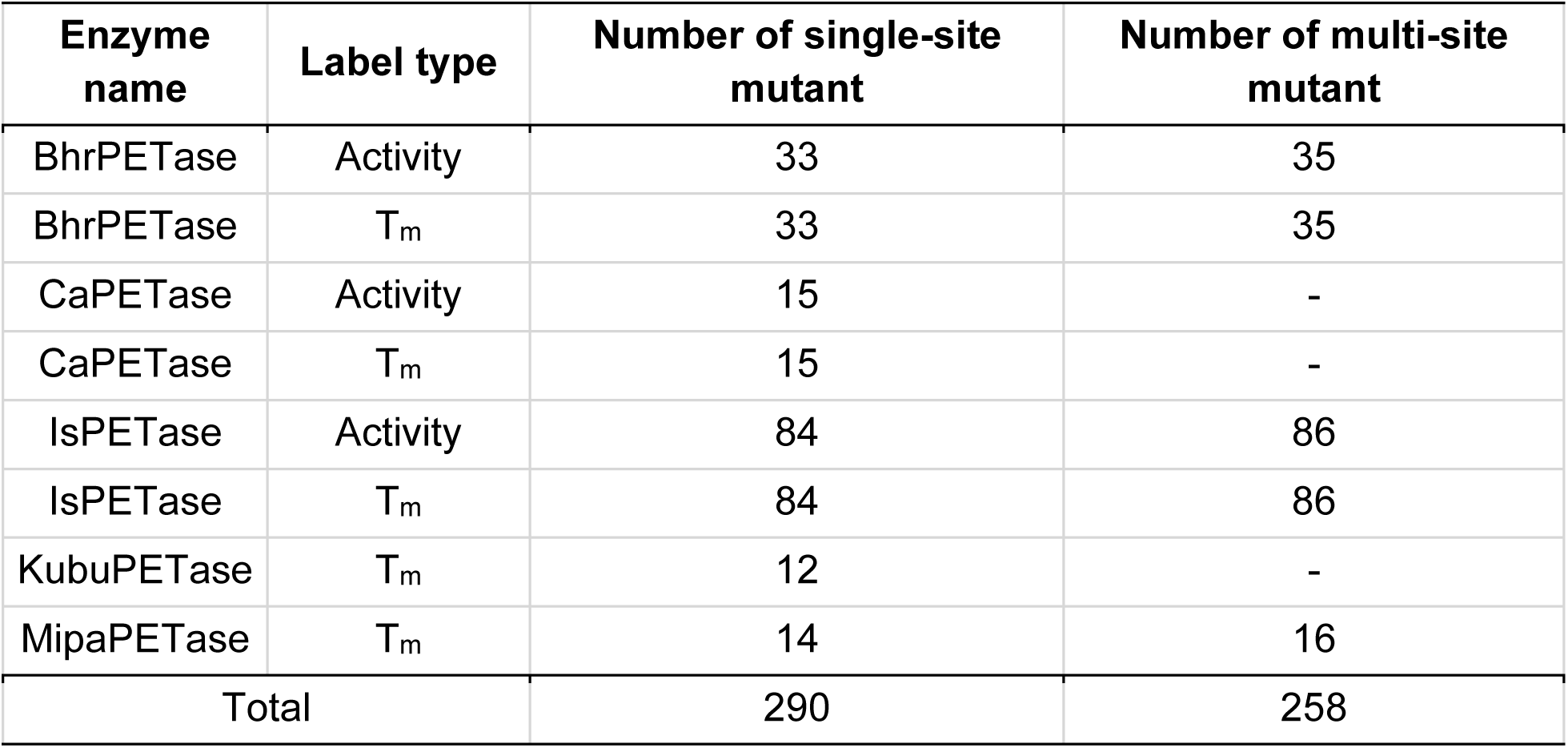
Summary statistics of the PET-Gym benchmark dataset.

**Supplementary Table 5.** Machine learning algorithms evaluated in PET-Gym (single-to-high).

| Abbrevia<br>tion | Full Name | Description |
| --- | --- | --- |
| SVR | Support Vector Regression | Regression using support vector machines with epsilon-insensitive loss |
| NuSVR | Nu Support Vector Regression | Similar to SVR but uses a parameter nu to control the number of support vectors |
| MLP | Multi-Layer Perceptron | Neural network with one hidden layers for regression |
| Lasso | Lasso Regression | Linear regression with L1 regularization (promotes sparsity) |
| Ridge | Ridge Regression | Linear regression with L2 regularization (prevents large coefficients) |
| ElasticNet | Elastic Net Regression | Linear regression combining L1 and L2 regularization |
| Bayesian Ridge | Bayesian Ridge Regression | Ridge regression with Bayesian inference for automatic regularization tuning |
| Huber | Huber Regressor | Robust linear regression less sensitive to outliers |
| KNN | K-Nearest Neighbors Regressor | Predicts based on the average of k nearest training samples |
| GPR | Gaussian Process Regression | Non-parametric probabilistic regression using kernel functions |
| KernelRidge | Kernel Ridge Regression | Ridge regression combined with kernel trick for non-linear patterns |

**Supplementary Table 6.** Amino acid sequences of M1-M32.

**Supplementary Table 7.**
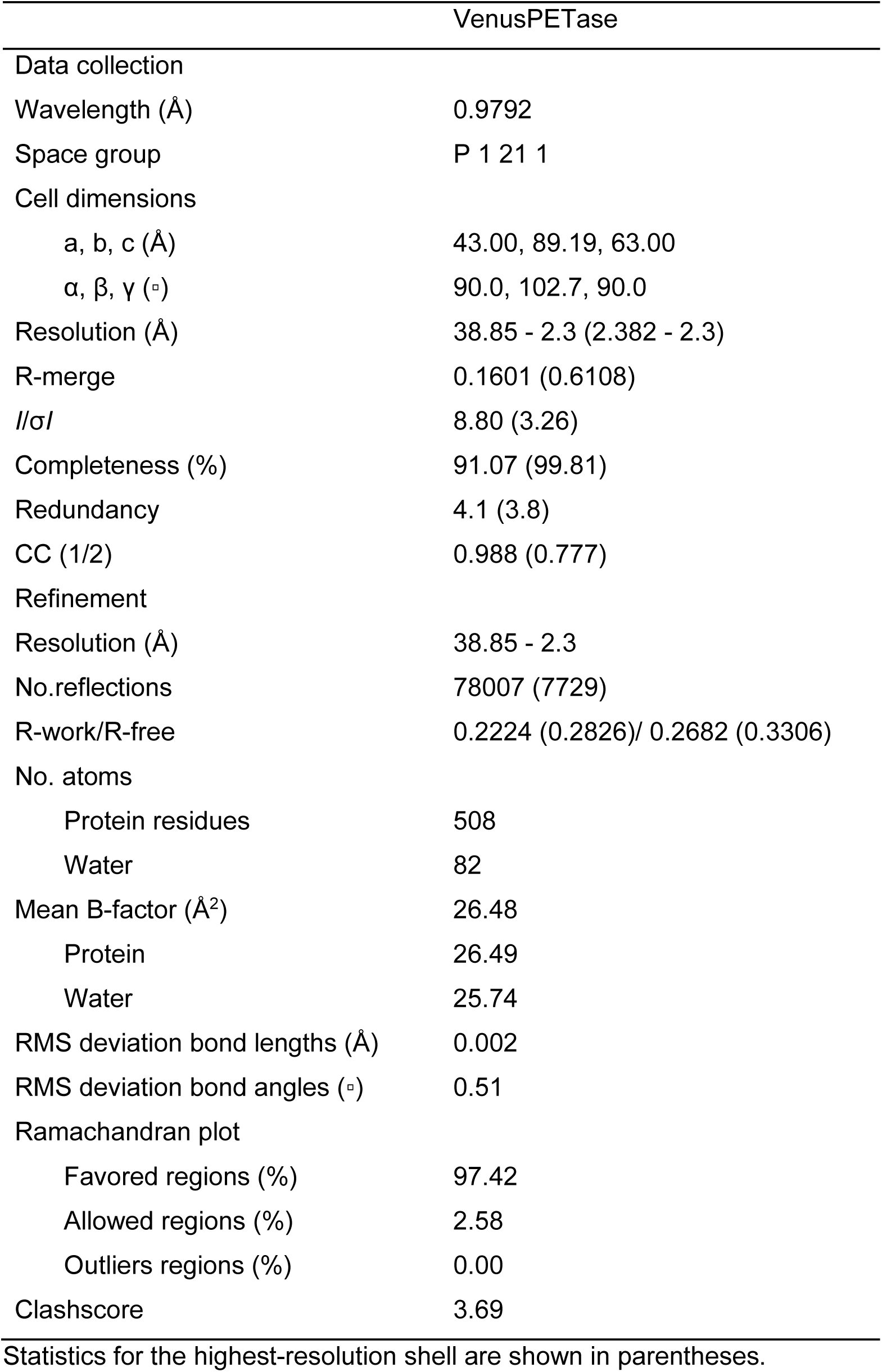
Data collection and refinement statistics.

|  | VenusPETase |
| --- | --- |
| Data collection |  |
| Wavelength (Å) | 0.9792 |
| Space group | P 1 21 1 |
| Cell dimensions |  |
| a, b, c (Å) | 43.00, 89.19, 63.00 |
| $\alpha$, $\beta$, $\gamma$ (°) | 90.0, 102.7, 90.0 |
| Resolution (Å) | 38.85 - 2.3 (2.382 - 2.3) |
| R-merge | 0.1601 (0.6108) |
| *I*/σ*I* | 8.80 (3.26) |
| Completeness (%) | 91.07 (99.81) |
| Redundancy | 4.1 (3.8) |
| CC (1/2) | 0.988 (0.777) |
| Refinement |  |
| Resolution (Å) | 38.85 - 2.3 |
| No.reflections | 78007 (7729) |
| R-work/R-free | 0.2224 (0.2826)/ 0.2682 (0.3306) |
| No. atoms |  |
| Protein residues | 508 |
| Water | 82 |
| Mean B-factor (Å<sup>2</sup>) | 26.48 |
| Protein | 26.49 |
| Water | 25.74 |
| RMS deviation bond lengths (Å) | 0.002 |
| RMS deviation bond angles (°) | 0.51 |
| Ramachandran plot |  |
| Favored regions (%) | 97.42 |
| Allowed regions (%) | 2.58 |
| Outliers regions (%) | 0.00 |
| Clashscore | 3.69 |

Statistics for the highest-resolution shell are shown in parentheses.

## References

1. R. Geyer, J. R. Jambeck, K. L. Law, Production, use, and fate of all plastics ever made. Science Advances 3, e1700782 (2017).

2. H. Lu et al., Machine learning-aided engineering of hydrolases for PET depolymerization. Nature 604, 662–667 (2022).

3. J. W. Cottom, E. Cook, C. A. Velis, A local-to-global emissions inventory of macroplastic pollution. Nature 633, 101--108 (2024).

4. J. Zhao et al., Microplastic fragmentation by rotifers in aquatic ecosystems contributes to global nanoplastic pollution. Nature nanotechnology 19, 406--414 (2024).

5. R. G. Santos, G. E. Machovsky-Capuska, R. Andrades, Plastic ingestion as an evolutionary trap: Toward a holistic understanding. Science 373, 56--60 (2021).

6. J. Chen et al., Global marine microbial diversity and its potential in bioprospecting. Nature 633, 371–379 (2024).

7. M. MacLeod, H. P. H. Arp, M. B. Tekman, A. Jahnke, The global threat from plastic pollution. Science 373, 61--65 (2021).

8. V. Nava et al., Plastic debris in lakes and reservoirs. Nature 619, 317–322 (2023).

9. J. R. Jambeck et al., Plastic waste inputs from land into the ocean. Science 347, 768–771 (2015).

10. E. J. Carpenter, K. L. Smith Jr, Plastics on the Sargasso Sea surface. Science 175, 1240--1241 (1972).

11. C. Ostle et al., The rise in ocean plastics evidenced from a 60-year time series. Nature communications 10, 1622 (2019).

12. S. B. Borrelle et al., Predicted growth in plastic waste exceeds efforts to mitigate plastic pollution. Science 369, 1515–1518 (2020).

13. W. W. Y. Lau et al., Evaluating scenarios toward zero plastic pollution. Science 369, 1455--1461 (2020).

14. N. Simon et al., A binding global agreement to address the life cycle of plastics. Science 373, 43--47 (2021).

15. J. M. Garcia, M. L. Robertson, The future of plastics recycling. Science 358, 870--872 (2017).

16. L. D. Ellis et al., Chemical and biological catalysis for plastics recycling and upcycling. Nature Catalysis 4, 539–556 (2021).

17. C. Jehanno et al., Critical advances and future opportunities in upcycling commodity polymers. Nature 603, 803--814 (2022).

18. N. W. Johnson et al., A biocompatible Lossen rearrangement in Escherichia coli. Nature Chemistry, 1--7 (2025).

19. F. Vidal et al., Designing a circular carbon and plastics economy for a sustainable future. Nature 626, 45–57 (2024).

20. V. Tournier et al., Enzymes’ Power for Plastics Degradation. Chemical Reviews 123, 5612–5701 (2023).

21. R. Wei, P. Westh, G. Weber, L. M. Blank, U. T. Bornscheuer, Standardization guidelines and future trends for PET hydrolase research. Nature Communications 16, 4684 (2025).

22. R. Wei et al., Possibilities and limitations of biotechnological plastic degradation and recycling. Nature Catalysis 3, 867–871 (2020).

23. W. Zimmermann, Polyester-degrading enzymes in a circular economy of plastics. Nature Reviews Bioengineering, 1--16 (2025).

24. S. Y. Choi et al., Sustainable production and degradation of plastics using microbes. Nature microbiology 8, 2253--2276 (2023).

25. S. Yoshida et al., A bacterium that degrades and assimilates poly(ethylene terephthalate). Science 351, 1196–1199 (2016).

26. Y. Cui et al., Computational Redesign of a PETase for Plastic Biodegradation under Ambient Condition by the GRAPE Strategy. ACS Catalysis 11, 1340–1350 (2021).

27. L. Shi et al., Complete Depolymerization of PET Wastes by an Evolved PET Hydrolase from Directed Evolution. Angew Chem Int Ed Engl 62, e202218390 (2023).

28. E. L. Bell et al., Directed evolution of an efficient and thermostable PET depolymerase. Nature Catalysis 5, 673–681 (2022).

29. H. F. Son et al., Rational protein engineering of thermo-stable PETase from Ideonella sakaiensis for highly efficient PET degradation. Acs Catalysis 9, 3519--3526 (2019).

30. R. Wei et al., Mechanism-based design of efficient PET hydrolases. ACS catalysis 12, 3382--3396 (2022).

31. T. B. Thomsen, C. J. Hunt, A. S. Meyer, Influence of substrate crystallinity and glass transition temperature on enzymatic degradation of polyethylene terephthalate (PET). New Biotechnology 69, 28–35 (2022).

32. B. C. Knott et al., Characterization and engineering of a two-enzyme system for plastics depolymerization. Proc Natl Acad Sci U S A 117, 25476–25485 (2020).

33. G. Arnal et al., Assessment of Four Engineered PET Degrading Enzymes Considering Large-Scale Industrial Applications. ACS Catalysis 13, 13156–13166 (2023).

34. P. Blázquez-Sánchez et al., Computational engineering of the polyester hydrolase PHL7 for efficient poly(ethylene terephthalate) degradation in biocatalytic recycling processes. Nature Communications 17, 4370 (2026).

35. H. Hong et al., Discovery and rational engineering of PET hydrolase with both mesophilic and thermophilic PET hydrolase properties. Nature communications 14, 4556 (2023).

36. B. Wu et al., Harnessing protein language model for structure-based discovery of highly efficient and robust PET hydrolases. Nature Communications 16, 6211 (2025).

37. V. Tournier et al., An engineered PET depolymerase to break down and recycle plastic bottles. Nature 580, 216–219 (2020).

38. H. Seo et al., Landscape profiling of PET depolymerases using a natural sequence cluster framework. Science 387, eadp5637 (2025).

39. L. Pfaff et al., Multiple Substrate Binding Mode-Guided Engineering of a Thermophilic PET Hydrolase. ACS Catal 12, 9790–9800 (2022).

40. H. F. Son et al., Rational Protein Engineering of Thermo-Stable PETase from Ideonella sakaiensis for Highly Efficient PET Degradation. ACS Catalysis 9, 3519–3526 (2019).

41. A. Bollinger et al., A Novel Polyester Hydrolase From the Marine Bacterium Pseudomonas aestusnigri – Structural and Functional Insights. Frontiers in Microbiology 11, (2020).

42. K. Dresler, J. van den Heuvel, R.-J. Müller, W.-D. Deckwer, Production of a recombinant polyester-cleaving hydrolase from Thermobifida fusca in Escherichia coli. Bioprocess and Biosystems Engineering 29, 169–183 (2006).

43. F. Kawai et al., A novel Ca²⁺-activated, thermostabilized polyesterase capable of hydrolyzing polyethylene terephthalate from Saccharomonospora viridis AHK190. Appl Microbiol Biotechnol 98, 10053–10064 (2014).

44. E. Herrero Acero et al., Enzymatic Surface Hydrolysis of PET: Effect of Structural Diversity on Kinetic Properties of Cutinases from Thermobifida. Macromolecules 44, 4632–4640 (2011).

45. R.-J. Müller, H. Schrader, J. Profe, K. Dresler, W.-D. Deckwer, Enzymatic Degradation of Poly(ethylene terephthalate): Rapid Hydrolyse using a Hydrolase from T. fusca. Macromolecular Rapid Communications 26, 1400–1405 (2005).

46. S. Sulaiman et al., Isolation of a Novel Cutinase Homolog with Polyethylene Terephthalate-Degrading Activity from Leaf-Branch Compost by Using a Metagenomic Approach. Applied and Environmental Microbiology 78, 1556–1562 (2012).

47. Y. Tan, B. Zhou, L. Zheng, G. Fan, L. Hong, Semantical and Topological Protein Encoding Toward Enhanced Bioactivity and Thermostability. bioRxiv, 2023--2012 (2023).

48. L. Zheng et al., Scalable and multiplexed recorders of gene regulation dynamics across weeks. Nature 652, 1038–1048 (2026).

49. Y. Cui et al., Computational redesign of a hydrolase for nearly complete PET depolymerization at industrially relevant high-solids loading. Nature Communications 15, 1417 (2024).

50. N. P. Murphy et al., Process innovations to enable viable enzymatic poly (ethylene terephthalate) recycling. Nature Chemical Engineering, 1--12 (2025).

51. S. W. Schubert et al., Relationships of crystallinity and reaction rates for enzymatic degradation of poly (ethylene terephthalate), PET. ChemSusChem 17, e202301752 (2024).

52. R. Vanella et al., Understanding activity-stability tradeoffs in biocatalysts by enzyme proximity sequencing. Nature Communications 15, 1807 (2024).

53. B. K. Shoichet, W. A. Baase, R. Kuroki, B. W. Matthews, A relationship between protein stability and protein function. Proceedings of the National Academy of Sciences 92, 452–456 (1995).

54. H. Yu, P. A. Dalby, Exploiting correlated molecular-dynamics networks to counteract enzyme activity–stability trade-off. Proceedings of the National Academy of Sciences 115, E12192–E12200 (2018).

55. A. Elnaggar, et al., Ankh: Optimized protein language model unlocks general-purpose modelling. arXiv:2301.06568, (2023).

56. D. M. Weinreich, N. F. Delaney, M. A. DePristo, D. L. Hartl, Darwinian Evolution Can Follow Only Very Few Mutational Paths to Fitter Proteins. Science 312, 111–114 (2006).

57. M. Wittmund, F. Cadet, M. D. Davari, Learning Epistasis and Residue Coevolution Patterns: Current Trends and Future Perspectives for Advancing Enzyme Engineering. ACS Catalysis 12, 14243–14263 (2022).

58. C. Fröhlich et al., Epistasis arises from shifting the rate-limiting step during enzyme evolution of a β-lactamase. Nature Catalysis 7, 499–509 (2024).

59. B. Eiamthong et al., Discovery and Genetic Code Expansion of a Polyethylene Terephthalate (PET) Hydrolase from the Human Saliva Metagenome for the Degradation and Bio-Functionalization of PET. Angewandte Chemie International Edition 61, e202203061 (2022).

60. A. Singh et al., Techno-economic, life-cycle, and socioeconomic impact analysis of enzymatic recycling of poly(ethylene terephthalate). Joule 5, 2479–2503 (2021).

61. J. Cao, H. Liang, W. Chen, X. Li, S. Fu, Sustainable recycling of polyester wastes using a coordinatively unsaturated Zn catalyst. Nature Communications, (2026).

62. J. Zhou, Z. Cui, R. Wei, W. Dong, M. Jiang, Interfacial catalysis in enzymatic PET plastic depolymerization. Trends in Chemistry 7, 175–185 (2025).

63. N. A. Tarazona et al., Rapid depolymerization of poly(ethylene terephthalate) thin films by a dual-enzyme system and its impact on material properties. Chem Catalysis 2, 3573–3589 (2022).

64. Y. Zhang et al., Glass Transition Temperature Determination of Poly(ethylene terephthalate) Thin Films Using Reflection−Absorption FTIR. Macromolecules 37, 2532–2537 (2004).

65. K. Shinotsuka, V. N. Bliznyuk, H. E. Assender, Near-surface crystallization of PET. Polymer 53, 5554–5559 (2012).

66. Y.-J. Liu et al., State-of-the-art advances in biotechnology for polyethylene terephthalate bio-depolymerization. Green Carbon, (2025).

67. Y.-H. V. Soong, M. J. Sobkowicz, D. Xie, Recent Advances in Biological Recycling of Polyethylene Terephthalate (PET) Plastic Wastes. Bioengineering 9, 98 (2022).

68. N. E. Wallace et al., The highly crystalline PET found in plastic water bottles does not support the growth of the PETase-producing bacterium Ideonella sakaiensis. Environmental Microbiology Reports 12, 578–582 (2020).

69. S. Bartwal, A. Matura, R. Kumar, H. Malhotra, Life cycle assessment of polyethylene terephthalate (PET): an end-of-life perspective. Green Materials, 1–17 (2025).

70. A. Patel et al., Melt Processing Pretreatment Effects on Enzymatic Depolymerization of Poly(ethylene terephthalate). ACS Sustainable Chemistry & Engineering 10, 13619–13628 (2022).

71. A. A. Modenbach, S. E. Nokes, Enzymatic hydrolysis of biomass at high-solids loadings – A review. Biomass and Bioenergy 56, 526–544 (2013).

72. M. D. de Dios Caputto, R. Navarro, J. L. Valentín, Á. Marcos-Fernández, Chemical upcycling of poly(ethylene terephthalate) waste: Moving to a circular model. Journal of Polymer Science 60, 3269–3283 (2022).

73. Z. Chen, H. Sun, W. Kong, L. Chen, W. Zuo, Closed-loop utilization of polyester in the textile industry. Green Chemistry 25, 4429–4437 (2023).

74. R. Wei et al., Biocatalytic Degradation Efficiency of Postconsumer Polyethylene Terephthalate Packaging Determined by Their Polymer Microstructures. Advanced Science 6, 1900491 (2019).

75. S. R. Eddy, Accelerated profile HMM searches. PLoS Computational Biology 7, e1002195 (2011).

76. Z. Lin et al., Evolutionary-scale prediction of atomic-level protein structure with a language model. Science 379, 1123–1130 (2023).

77. UniProt: the Universal Protein Knowledgebase in 2025. Nucleic Acids Res 53, D609-d617 (2025).

78. K. K. Yang, N. Zanichelli, H. Yeh, Masked inverse folding with sequence transfer for protein representation learning. *Protein Engineering*, Design and Selection 36, gzad015 (2023).

79. J. Meier et al., Language models enable zero-shot prediction of the effects of mutations on protein function. Advances in Neural Information Processing Systems 34, 29287--29303 (2021).

80. Z. Zhang et al., 2024.

81. Y. Tan, R. Wang, B. Wu, L. Hong, B. Zhou, From high-throughput evaluation to wet-lab studies: advancing mutation effect prediction with a retrieval-enhanced model. Bioinformatics 41, i401 (2025).

82. J. Su et al., Democratizing protein language model training, sharing and collaboration. Nature Biotechnology, (2025).

83. B. Chen et al., xTrimoPGLM: unified 100-billion-parameter pretrained transformer for deciphering the language of proteins. Nature Methods 22, 1028–1039 (2025).

84. M. J. Frisch, et al. (2016).

85. P. J. Stephens, F. J. Devlin, C. F. Chabalowski, M. J. Frisch, Ab Initio Calculation of Vibrational Absorption and Circular Dichroism Spectra Using Density Functional Force Fields. Journal of Physical Chemistry 98, 11623–11627 (1994).

86. F. Weigend, Accurate Coulomb-fitting basis sets for H to Rn. Physical Chemistry Chemical Physics 8, 1057–1065 (2006).

87. S. Grimme, S. Ehrlich, L. Goerigk, Effect of the damping function in dispersion corrected density functional theory. Journal of Computational Chemistry 32, 1456–1465 (2011).

88. M. Schauperl et al., Non-bonded force field model with advanced restrained electrostatic potential charges (RESP2). Communications Chemistry 3, 44 (2020).

89. T. Lu, F. Chen, Multiwfn: a multifunctional wavefunction analyzer. Journal of Computational Chemistry 33, 580–592 (2012).

90. J. Zhang, T. Lu, Efficient evaluation of electrostatic potential with computerized optimized code. Physical Chemistry Chemical Physics 23, 20323–20328 (2021).

91. S. Páll et al., Heterogeneous parallelization and acceleration of molecular dynamics simulations in GROMACS. The Journal of Chemical Physics 153, 134110 (2020).

92. J. Wang, R. M. Wolf, J. W. Caldwell, P. A. Kollman, D. A. Case, Development and testing of a general amber force field. Journal of Computational Chemistry 25, 1157–1174 (2004).

93. G. Bussi, D. Donadio, M. Parrinello, Canonical sampling through velocity rescaling. Journal of Chemical Physics 126, 014101 (2007).

94. W. L. Jorgensen, J. Chandrasekhar, J. D. Madura, R. W. Impey, M. L. Klein, Comparison of simple potential functions for simulating liquid water. Journal of Chemical Physics 79, 926–935 (1983).

95. M. Bernetti, G. Bussi, Pressure control using stochastic cell rescaling. The Journal of Chemical Physics 153, 114107 (2020).

96. U. Essmann et al., A smooth particle mesh Ewald method. Journal of Chemical Physics 103, 8577–8593 (1995).

97. T. D. Kühne et al., CP2K: An electronic structure and molecular dynamics software package - Quickstep: Efficient and accurate electronic structure calculations. The Journal of Chemical Physics 152, 194103 (2020).

98. C. Bai, R. J. Spontak, C. C. Koch, C. K. Saw, C. M. Balik, Structural changes in poly(ethylene terephthalate) induced by mechanical milling. Polymer 41, 7147–7157 (2000).

99. J. P. Perdew, K. Burke, M. Ernzerhof, Generalized Gradient Approximation Made Simple. Physical Review Letters 77, 3865–3868 (1996).

100. J. VandeVondele, J. Hutter, Gaussian basis sets for accurate calculations on molecular systems in gas and condensed phases. The Journal of Chemical Physics 127, 114105 (2007).

101. T. Lu, A comprehensive electron wavefunction analysis toolbox for chemists, Multiwfn. The Journal of Chemical Physics 161, (2024).

102. T. Lu. (2024).

103. W. L. Delano, The PyMol Molecular Graphics System. Proteins Structure Function \& Bioinformatics 30, 442–454 (2002).

104. W. Humphrey, A. Dalke, K. Schulten, VMD: Visual molecular dynamics. Journal of Molecular Graphics 14, 33–38 (1996).

105. J. A. Maier et al., ff14SB: Improving the Accuracy of Protein Side Chain and Backbone Parameters from ff99SB. Journal of Chemical Theory and Computation 11, 3696–3713 (2015).

106. B. Hess, H. Bekker, H. J. C. Berendsen, J. G. E. M. Fraaije, LINCS: A linear constraint solver for molecular simulations. Journal of Computational Chemistry 18, 1463–1472 (1997).

107. N. Michaud-Agrawal, E. J. Denning, T. B. Woolf, O. Beckstein, MDAnalysis: A toolkit for the analysis of molecular dynamics simulations. Journal of Computational Chemistry 32, 2319–2327 (2011).

108. Y. Huang et al., DSDP: A Blind Docking Strategy Accelerated by GPUs. Journal of Chemical Information and Modeling 63, 4355–4363 (2023).

109. J. Zhang, T. Gu, C. Li, W. Qi, s_mmpbsa: A Lite and Cross-Platform MM-PBSA Program. Molecules 31, 1683 (2026).

110. G. Fiorin, M. L. Klein, J. Hénin, Using collective variables to drive molecular dynamics simulations. Molecular Physics 111, 3345–3362 (2013).

111. S. Grimme, C. Bannwarth, P. Shushkov, A Robust and Accurate Tight-Binding Quantum Chemical Method for Structures, Vibrational Frequencies, and Noncovalent Interactions of Large Molecular Systems Parametrized for All spd-Block Elements (Z = 1–86). J Chem Theory Comput 13, 1989-2009 (2017).

112. K. Huynh, C. L. Partch, Analysis of Protein Stability and Ligand Interactions by Thermal Shift Assay. Current Protocols in Protein Science 79, 28.29.21–28.29.14 (2015).

113. L. Shi et al., Complete depolymerization of PET wastes by an evolved PET hydrolase from directed evolution. Angewandte Chemie 135, e202218390 (2023).

114. W. Kabsch, Integration, scaling, space-group assignment and post-refinement. Acta Crystallographica Section D Biological Crystallography 66, 133–144 (2010).

115. P. Evans, G. Murshudov, How good are my data and what is the resolution? Acta Crystallographica Section D: Biological Crystallography 69, 1204 - 1214 (2013).

116. A. J. McCoy et al., Phaser crystallographic software. Journal of Applied Crystallography 40, (2007).

117. P. Emsley, K. Cowtan, Coot: model-building tools for molecular graphics. Acta Crystallogr D Biol Crystallogr 60, 2126–2132 (2004).

118. P. V. Afonine et al., Towards automated crystallographic structure refinement with phenix.refine. Acta Crystallographica 68, 352–367 (2012).

119. C. J. Williams et al., MolProbity: More and better reference data for improved all-atom structure validation. Protein Science 27, 293–315 (2018).

